# LDB1 regulates gene expression and chromatin structure in pluripotency and lineage differentiation

**DOI:** 10.1101/2025.01.17.633568

**Authors:** HeungSun Kwon, Juhyun Kim, Lecong Zhou, Ann Dean

## Abstract

Chromatin organization is a pivotal factor in stem cell pluripotency and differentiation. However, the role of enhancer looping protein LDB1 in stem cells has not been explored. We generated *Ldb1(−/−)* embryonic stem cells (ESC) using CRISPR/Cas9 editing and observed a reduction in key stem cell factors SOX2 and KLF4 upon LDB1 loss. Embryoid bodies (EB) derived from *Ldb1(−/−)* ESC displayed reduced expression of lineage-specific markers and impaired ability to undergo terminal differentiation to erythroblasts. Differential gene expression, including of the *Lin28*-mediated self-renewal pathway genes, was observed between WT and *Ldb1(−/−)* ESC and EB but was most pronounced after differentiation to erythroblasts. LDB1 occupied super enhancers, including those of pluripotency genes, in ESC together with pluripotency factors. LDB1 loss resulted in globally decreased chromatin accessibility in ESC and EB. Conditional LDB1-deficient mice displayed reduced hematopoietic stem cell markers on bone marrow cells, and dysregulation of the *Lin28* pathway. Thus, LDB1 function is critical for ESC and EB development and becomes progressively more important during differentiation to erythroblasts.

## INTRODUCTION

Chromatin organization plays a vital role in development and cell lineage choice (1,2). At the same time, precise gene expression control is crucial for proper cell fate determination and tissue formation. This transcriptional specificity is orchestrated by the activation and decommissioning of enhancers and their long-range interactions within the context of overall genome folding (3–6). Transcriptional enhancers have a key role by determining which genes are active and which are silent (7).

LIM domain-binding 1 (LDB1 is part of multi-protein complexes that drive interaction, or looping, between enhancers and target genes in erythroid cells and in diverse other tissue types by partnering with specific LIM-homeodomain and LIM-only factors to access chromatin (8–17). LDB1 enhancer looping is instructive for gene expression, a mechanism that becomes more prevalent as calls transition from fate specification to tissue differentiation (18,19). *Ldb1(−/−)* mouse embryos die before E8.5 with evidence of malformations in multiple tissue types, indicating that LDB1 is essential for normal development (20). LDB1 complexes are implicated in various cellular processes beyond chromatin looping, including chromatin remodeling and direct participation in transcriptional regulation (21–24). In the hematopoietic lineage, LDB1 complexes underlie developmental progression (25–27). The role of LDB1 in embryonic stem cells and in lineage choice has not been explored.

Stem cell proliferation and differentiation are tightly controlled processes involving a discrete number of transcription factors (28–30). SOX2, OCT4, and KLF4 are key proteins that maintain pluripotency and self-renewal of embryonic stem cells (31). These pluripotency factors bind to enhancers and large super enhancers together with the Mediator coactivator complex in embryonic stem cells and function in long range enhancer-promoter interactions in these cells (32–34). Pluripotency factors have also been implicated in modulating chromatin accessibility dynamics and transcriptional activity during induced pluripotent stem cell reprogramming (35,36). Moreover, KLF4 interacts with OCT4 and SOX2 and is thought to have a causal role in enhancer looping in embryonic stem cells (37–39).

Pluripotency factors further engage in more extensive regulatory interactions in embryonic stem cells, including with NODAL and WNT networks (40,41). LIN28 is another crucial regulator of stem cell activity and development that is involved in both self-renewal and differentiation (42). The interplay between LIN28 and SOX2, OCT4, and KLFs is important for maintaining stem cell pluripotency, regulating lineage-specific differentiation and induced pluripotent stem cell reprogramming (43–45). The LIN28b-*Let-7*-HMGA pathway critically underlies the self-renewal potential of fetal hematopoietic stem cells (46).

Here we provide evidence for the role of LDB1 in the proliferation and development of embryonic stem cells, embryoid bodies and erythroblasts differentiated from them. Loss of LDB1 dysregulates pluripotency genes as well as interacting networks involving NODAL, WNT and LIN28 signaling by both direct and indirect mechanisms. The LDB1 enhancer landscape is modulated during embryonic stem cell differentiation and becomes extensive upon differentiation of stem cells to erythroblasts. Our findings provide insight into the molecular mechanisms that underlie embryonic stem cell proliferation and differentiation, with LDB1 emerging as an important player in orchestrating these processes, including through its involvement in super enhancer function.

## MATERIALS AND METHODS

### ESC culture and EB formation

Wild-type ESC (ES-E14G2a) were obtained from ATCC (Manassas, VA). ESC were maintained and differentiated as described, with modifications (47). Undifferentiated ESC were cultured on 0.1% gelatin-coated plates in EmbryoMax DMEM (MilliporeSigma, SLM-220-M supplemented with 10% fetal bovine serum (MilliporeSigma), 2-β-mercaptoethanol (MilliporeSigma, ES-007-E), GlutaMax, penicillin/streptomycin, non-essential amino acids, and ESGRO-2i (MilliporeSigma, cat# SF016-200. To generate EB, cultures were adjusted to 1 × 10^5 cells/ml and incubated in EB medium (STEMCELL TECH, Aggrewell EB formation medium) for 3 to 5 days in Aggrewell plates (STEMCELL TECH, 34421). After 4 days, EB were pooled and grown in suspension in non-adherent plates, with media changed daily. EB were collected on days 3 and 5 for analysis and imaged using a microscope (model # RVL-100-G, ECHO) at 40x magnification. EB size was determined using ImageJ, and statistical significance of differences in size was determined using the Wilcoxon test in R.

### Primitive erythrocyte differentiation in vitro

EBs were differentiated as described (Cai et al., 2020). Briefly, EBs grown in AggreWell™ plates for 3 days were dissociated into single cells and cultured for 8 days with replacement of media on day 4. Cultures were then sorted into CD71+ /Ter119+ populations by FACS for analysis. Dead cells were excluded using DAPI (ThermoFisher, D1306).

### Conditional Ldb1 knock out mice

Mouse protocols were approved by the NIDDK Animal Care and Use Committee in accordance with AALAC specifications. *Ldb1*^flox/flox^ mice (kind gift of Dr. Yangu Zhou) were bred to *Mx-Cre* C57BL/6 mice (Jackson Laboratories) to obtain conditional *Ldb1* knock out animals. To delete *Ldb1*, 8-to 12-week-old male mice wild-type (WT) and *Mx-Cre; Ldb1*^flox/flox^ were subjected to intraperitoneal injections of synthetic double-stranded RNA (dsRNA) PolyI:PolyC or PIPC (1 mg/mL in PBS) (or PBS control) 3 times in 1 week to induce *in vivo* production of IFN-a/h, subsequently triggering the activation of the *Mx-Cre* transgene (48). Mice were euthanized after 14 days. Liver tissues were collected and utilized to confirm the successful deletion of *Ldb1*.

### Ldb1(−/−) ESC

CRISPR/Cas9 genome editing was used to generate *Ldb1*(−/−) ES cells. CRISPR gRNAs were designed using the http://crispr.mit.edu/ wEBite (see Supplemental Table S1 for *gRNA* sequences). *Ldb1*-targeting gRNAs were cloned into the CRISPR-Cas9 and gRNA expression vector pSpCas9(BB)-2A-GFP (PX458) (Addgene, cat #48138). ES cells were transfected with Mouse ES Cell Nucleofector® Kit (Lonza, cat # VPH-1001) according to the manufacturer’s instructions. Fluorescent cells were sorted 48 hours later and plated at limiting dilution to isolate single clones harboring the LDB1 gene knockout. See Table S1 for gRNAs.

### Tet-inducible LDB1 expression plasmid

A Tet-inducible LDB1 expression plasmid was generated using the pLIX_403 vector (Addgene, 41395). The LDB1 gene was inserted into the destination vector pLIX_403 using the gateway cloning system (pENTRY-LDB1). The plasmid was transfected into HEK293 cells using Lipo300 transfection reagent (ThermoFisher, L3000075). Transfected cells were treated with doxycycline (1 mg/ml) to activate the Tet-inducible promoter in the pLIX_403 vector.

### FACS analysis

Flow cytometry was performed using a BD Accuri C6 Flow Cytometer (BD Biosciences) and data was analyzed with FlowJo software (Tree Star, Inc.). Cells were labeled with TER119-PE and CD71-FITC antibodies and processed using the MACS® MiniSampler of the autoMACS® Pro Separator (Miltenyi Biotec) according to the manufacturer’s protocol. Cell enrichment was performed using program positive selection with TER119 bead (Miltenyi Biotec, 130-049-901). Samples were consecutively processed after mixing to resuspend cells. See Table S2 for antibodies.

### Lentiviral preparation

5 x 10^6^ 293T cells were seeded in 15 cm culture dishes with 10 mL DMEM supplemented with 10% fetal bovine serum, 1% HEPES, 1% non-essential amino acids (NEAA), 1% penicillin/streptomycin, and 1% L-Glutamine. Fresh media was added after 24 hours and co-transfection performed with 8 µg of a lentiviral expression plasmid and 6 µg psPAX2 and 4 µg pMD2.G using polyethyleneimine. 12-16 hours post transfection, the media was replaced with 5mL DMEM supplemented with 5% FBS, 1% HEPES, and 1% NEAA. Viral supernatants were collected 48 and 72 hours post transfection and filtered through 0.45 µm filters. Viral supernatants were used fresh to infect target cells or concentrated by ultracentrifugation and snap frozen, stored at –80°C for future use.

### EdU incorporation

5-ethynyl-uridine (EdU)-labeling was performed using Click-iT EdU Imaging Kits (Invitrogen, C10646), following the manufacturer’s instructions. Briefly, 1×10^5^ ESC were seeded per well in 8-well glass chamber gelatin coated slides. EdU (10 nM) was added for 24 h. After labeling, cells were washed two to three times with PBS followed by addition of tissue culture media. For immediate analysis, washes were omitted, and cells were permeabilized and fixed by standard formaldehyde fixation protocol.

### Alkaline phosphatase staining

Alkaline phosphatase live cell staining was performed according to the manufacturer’s instructions (ThermoFisher, A14353) in basal DMEM/F-12 media. Images were captured within 30–60 min post-staining and the colonies exhibiting the strongest fluorescence were selected and expanded. After visualization, the basal media was replaced by fresh hESC growth media and the selected colonies were either manually picked or returned to the normal culture conditions.

### qRT-PCR

Total RNA (1 μg) treated with DNase I was reverse transcribed using Superscript III according to the manufacturer’s protocol (Invitrogen). The cDNA was diluted to 200 μl and 2 μl was used for real-time quantitative PCR (RT-qPCR) in a reaction volume of either 10 μl or 25 μl, employing SYBR chemistry. The miR-Let7a real-time PCR analysis was conducted utilizing TaqMan primers and labeled probe sets (ThermoFisher, 4427975. The primer snoRNA202 (ThermoFisher, 4427975, served as a control. At least three independent RNA preparations were analyzed and data normalized to *Gapdh*. Error bars indicate ±SEM, and the data were analyzed using the 2^−ΔΔCT^ relative quantitation method. See Table S1 for primers.

### Western Blotting

Cells were lysed in RIPA buffer and samples prepared for Western blot by boiling in 1× SDS loading buffer for 5 min, followed by electrophoresis on 10% SDS-PAGE gels. Proteins were transferred to a PVDF membrane followed by blocking with 5% nonfat milk for 1 hour and incubation with primary antibodies overnight in a cold room with shaking. Membranes were washed with TBST (50 mM Tris pH 8.0, 150 mM NaCl, 0.1% Tween 20) 3 times and HRP-conjugated incubated with secondary antibodies for 1 hour at room temperature. Membranes were washed 3 times TBST and developed with Western Bright ECL HRP substrate (ThermoFisher, 32209). See Table S2 for antibodies.

### Immunofluorescence

ESC were cultured on gelatin coated glass coverslips in 12-well plates overnight to facilitate cell adhesion. Cells were fixed with 4% paraformaldehyde (Electron Microscopy Sciences) and washed twice with PBS (Invitrogen). Slides were treated with PBS containing 0.2% Triton X-100 and 3% BSA fraction (Sigma) for 45 minutes at room temperature and incubated with primary antibodies overnight at 4 °C. Slides were washed with PBS containing 0.1% BSA and then incubated with corresponding secondary antibodies for 60 minutes at room temperature. Immunofluorescence studies were conducted on a Zeiss LSM780 confocal microscope equipped with a 40X objective using DAPI staining to visualize nuclei. Acquired images were further processed using Fiji software v2.1.0/1.53c. See Table S2 for antibodies.

### Giemsa staining

Cells were fixed on slides in methanol for 5-7 minutes, air dried and incubated in Giemsa Stain solution (Sigma, GS500) diluted 1:20 with deionized water) for 15-60 minutes. Slides were then rinsed in deionized water and air dried before visualization under a microscope.

### Immunoprecipitation

Immunoprecipitation was performed using the Dynabead Protein G/A protocol (ThermoFisher). Briefly, 3.5 × 10^5^ cells/well were disbursed in 6-well plates. After 48 hours, cells were washed twice with PBS and lysed for 1 hour on ice with 1 ml of RIPA buffer (Sigma) supplemented with 1% protease inhibitors (Sigma). Lysed cells were centrifuged 10 minutes at 13,000 × g, and the supernatant from one well was mixed with 30 μl of Dynabeads Protein G/A beads (ThermoFisher) coupled with specific antibodies. The cell lysate and the antibody coupled Dynabeads were incubated together at 4 °C on a rocker platform overnight. Dynabeads were centrifuged for 1 minute at 1000 × g, and the beads washed three times with 1 ml of cold lysis. Antibody-coupled beads were suspended in 60 μl of electrophoresis buffer, heated to 95 °C for 10 minutes and centrifuged for 1 minute at 1000 × g and the supernatant was used for Western blot analysis. See Table S2 for antibodies.

### RNA sequencing

RNA-seq libraries were prepared from 1ug of total RNA from ESC, EB or differentiated cells using NEBNext Ultra TM RNA library Prep Kit for Illumina following manufacturer’s recommendations (NEB, Cat #7770). Library quality was validated with an Agilent Bioanalyzer 2100. Fifty base paired-end reads were performed with a NovaSeq 6000 instrument (Illumina).

### ChIPmentation

ChIPmentation was performed essentially as described (49). Briefly, 1×10^7^ mESC (ES-E14) cells were fixed with 2mM DSG followed by 1% formaldehyde, sonicated and immunoprecipitated with LDB1 antibody. Samples were bound to Protein A/G magnetic beads (Pierce™ ChIP-grade Protein A/G Magnetic Beads 26162) and tagmentated with 1ul of tagmentation enzyme (Illumina, Nextera XT DNA Library Preparation Kit, cat # FC-131–1024) for 10 min at 37°C. The beads were washed and the eluate de-crosslinked in ChIP elution buffer with Proteinase K for 1 h at 55°C and incubated at 65◦C overnight. DNA was amplified using Nextera custom primers and purified and size selected using SRI beads. Library quality was validated using a High Sensitivity DNA Chip in the Agilent Bioanalyzer 2100. Libraries were sequenced on a NovaSeq 6000 instrument (Illumina) by the Sequencing and Genomics Core Facility of NHLBI. See Table S1 for primers.

### ATAC seq

ATAC-seq was performed with the ATAC-Seq Kit (Active Motif, 53150), according to manufacturer’s protocol. Briefly, 50,000 cells were washed twice with PBS and incubated with 50 μl of tagmentation mix, consisting of 25 μl of 2x TD buffer, 2.5 μl of TDE1 enzyme, 0.5 μl of 1% digitonin, 16.4 μl of 1x PBS, and 5.6 μl of nuclease-free water, for 30 minutes at 37°C. After tagmentation, DNA was purified for PCR amplification using custom-made barcoded primers. Library quality was validated using a High Sensitivity DNA Chip in the Agilent Bioanalyzer 2100. Libraries were sequenced on a NovaSeq 6000 instrument (Illumina) by the NIDDK Genomics Core.

### CUT and Tag

CUT&Tag library preparation was performed with the CUT&Tag-IT™ Assay Kit (Active Motif, 53170) according to manufacturer’s protocol using 50,000 ESC or EB cells. 0.1ng of E. coli Spike-in DNA (EpiCypher 18-1401) was added to each sample at the pAG-Tn5 step (EpiCypher 15-1117). Universal i5 primers and uniquely barcoded i7 primers were added with 14 cycles of PCR. Libraries were sequenced by the NIDDK Genomics Core Facility with an Illumina NovaSeq 6000p= instrument. See Table S2 for antibodies.

### Mouse peripheral blood mononuclear cells (PBMC)

Whole blood was diluted with an equal volume of Dulbecco’s Phosphate-Buffered Saline (D.PBS) lacking Ca2+ and Mg2+ and layered onto an equal volume of Ficoll-Paque Plus (Cytiva, 17144002) in a 50 ml Falcon tube. Centrifugation was performed with a swinging bucket rotor at 400 × g for 30 minutes at room temperature (RT), with centrifuge brake disengaged. PBMCs, located at Ficoll-Paque Plus/plasma interface, were transferred to a fresh 50 ml Falcon tube. PBMCs were then subjected to two washes with D.PBS, each followed by centrifugation at 300 × g for 10 minutes.

### Isolation of mouse bone marrow

Bone marrow was flushed from tibia and femurs and plated in cell culture dishes in DMEM growth medium with 1 g/L D-glucose and 10% fetal bovine serum.

### Quantitation and statistical analysis

All experimental results were from 3 biological replicates. All quantified data were statistically analyzed and presented as mean ± SEM. Unless otherwise stated, one-way ANOVA was used to calculate statistical significance with P values as detailed in the text and figure legends. Additional details of data analysis appear in Supplemental Methods.

## RESULTS

### LDB1 deficiency downregulates pluripotency factors in embryonic stem cells

To gain insight into the role of LDB1 during development, we used CRISPR/Cas9 to generate *Ldb1(−/−)* embryonic stem cells (ESC) (Fig. S1A-C). Immunofluorescent staining revealed co-expression and co-localization of LDB1 and the pluripotency marker SSEA1 in WT ESC but not in *Ldb1(−/−)* cells (Fig. 1A). The absence of LDB1 protein in *Ldb1(−/−)* ES cells was confirmed by western blot (Fig. 1B). LDB1 loss resulted in altered stemness, as evidenced by increased alkaline phosphatase staining in *Ldb1(−/−)* cells (Fig. 1C). There was increased proliferation of ESC as measured by incorporation of EdU (Fig. 1D).

**Figure. 1.**
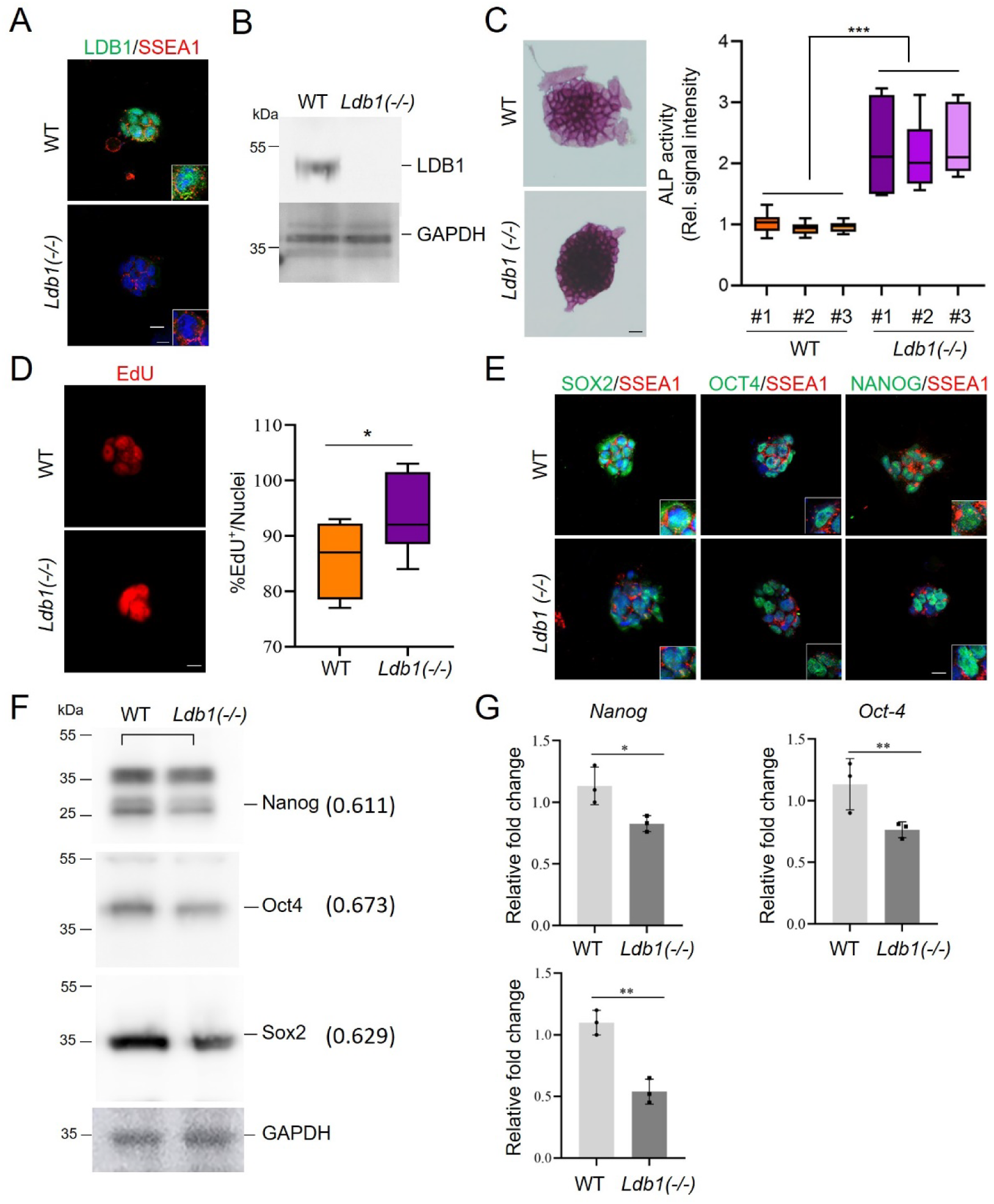
*Ldb1* deficiency impairs ESC proliferation and development (A) Immunofluorescent staining for LDB1 and SSEA1 in *Ldb1(−/−)* ESC. (B) Western blot of LDB1 in WT and *Ldb1(−/−)* ESC. (C) Alkaline phosphatase staining of *Ldb1(−/−)* and WT ESC. Quantitation (right) of alkaline phosphatase activity. N=10 for each of three independent clones. (D) 5-ethynyl-2’-deoxyuridine (EdU) incorporation in *Ldb1(−/−)* and WT ESC. Quantitation (right) of proliferation N=150. (E) Immunofluorescent staining of SOX2, OCT4, and NANOG, and stem cell surface marker SSEA1 in *Ldb1(−/−)* and WT ESC. (F) Western blot of key transcription factors involved in pluripotency and development in *Ldb1(−/−)* and WT ESC. (G) RT-qPCR of mRNA expression of *Nanog*, *Oct4*, and *Sox2* in *Ldb1(−/−)* ESC and WT ESC. *P<0.05, \*\**P* <0.01 and ***P<0.001 by two-tailed Student *t*-test; data shown as mean ±SEM. Scale bars, 5, 10 μm.

We next examined the effect of LDB1 loss on pluripotency factors SOX2, OCT4, and Nanog. Immunofluorescent staining, western blot, and mRNA analyses revealed reduction in the expression and protein levels of these factors in *Ldb1(−/−)* cells compared to WT ES cells (Fig. 1E-G). We confirmed and quantitated the decrease in SOX2 using immunofluorescent staining and established reduction of pluripotency factor KLF4 upon LDB1 loss (Fig. S1D, E). Additionally, the population of SOX2, OCT4 and SSEA1 expressing cells was dramatically reduced in *Ldb1(−/−)* ESC according to FACS analysis (Fig. S1 F-H). Thus, the loss of LDB1 significantly impacted the proliferation and stemness of ESC associated with reduction in the abundance of OCT4, SOX2, Nanog and KLF4.

### Impaired embryoid body formation and erythroblast development in *Ldb1(−/−)* ESC

To follow the influence of LDB1 during development, we differentiated *Ldb1(−/−)* and control ESC to embryoid bodies (EB) over 3 days of culture (Fig. 2A). We performed immunofluorescent staining and western blot analyses and documented formation of EB with no evidence of LDB1 protein (Fig. 2B, C). To quantitatively examine EB size, we seeded ESC on Aggrewell microplates. After 3 days of culture with EB medium, the *Ldb1(−/−)* EB exhibited a smaller size compared to WT EB (Fig. 2D). To assess the differentiation potential of *Ldb1(−/−)* EB, we examined the expression of three lineage-specific markers: *Gata1* (ectoderm), *Brachyury* (mesoderm), and *Gata-4* (endoderm). There was a significant reduction in the expression of these markers in *Ldb1(−/−)* EB, indicating impaired differentiation (Fig. 2E).

**Figure. 2.**
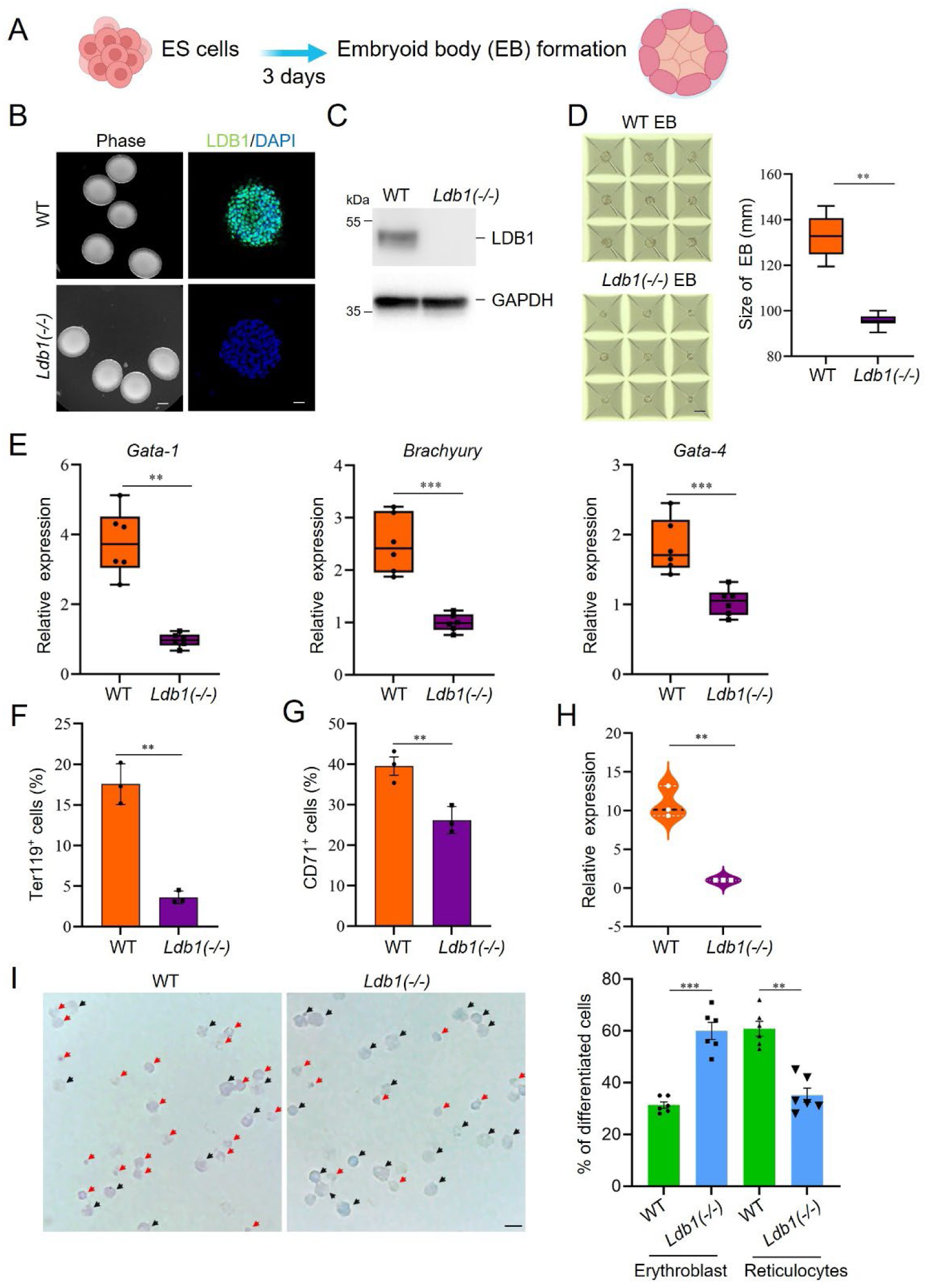
Impairment of differentiation due to *Ldb1* deficiency. (A) Protocol to generate embryoid bodies (EB) from ESC. (B) Phase contrast image and immunostaining of ESC for LDB1 after 3-day culture. Scale bars: 1 mm, 20 µm. (C) Immunoblot of LDB1 in WT and *Ldb1(−/−)* EB. (D) ESC were grown on Aggrewell plates for 3 days to form EB. Quantitation of EB size shown on right. (E) RT-qPCR of lineage markers in WT and *Ldb1(−/−)* EB. (F) Flow cytometry of TER119 on differentiated cells derived from WT and *Ldb1(−/−)* EB cells. (G) Flow cytometry of CD71 on differentiated cells derived from WT and *Ldb1(−/−)* EB. (H) RT-qPCR of β-globin in WT and *Ldb1(−/−)* after 8 days of differentiation from EB formation. (I) Giemsa staining of differentiated cells derived from WT and *Ldb1(−/−)* ESC. Blue arrows, erythroblasts; red arrows, reticulocytes. Scale bar: 50 µm. Quantitation of erythroblast and reticulocyte populations shown on the right (n=30). Data are mean ± SEM, n=3 biological replicates unless noted. Statistical significance determined by two-tailed Student’s t-test. * p<0.05, ** p<0.01, *** p<0.001.

To examine the effect of LDB1 loss on a fully differentiated cell lineage, we cultured dispersed EB in erythroid differentiation medium (26). To validate the efficacy of the differentiation protocol for WT EB, we examined the expression of β-globin, a well-established indicator of erythroid differentiation. The mRNA level of β-globin exhibited a progressive increase from day 8 to day 12, and protein expression was detectable at 12 days of incubation (Fig. S2A, B). Additionally, to confirm successful differentiation, we conducted FACS analysis of WT differentiated EB cells. The results demonstrated that approximately 41% and 45% of cells expressed the erythroid cell surface markers CD71 and Ter119, respectively, following 12 days of differentiation (Fig. S2C, D).

We next compared cells derived from WT and *Ldb1(−/−)* EB after differentiation in erythroid medium, documenting reduced numbers of cells expressing representative erythroid markers Ter119 and CD71 after LDB1 loss (Fig. 2F, G). Additionally, we found significant reduction of β-globin expression in *Ldb1(−/−)* differentiated cells (Fig. 2H). Moreover, upon Giemsa staining, we noted increased erythroblasts and decreased reticulocytes after loss of LDB1, consistent with delayed differentiation (Fig. 2I). In summary, the loss of LDB1 in ESC impaired or delayed EB formation and impaired erythroid differentiation of EB-derived cells.

### Differential gene expression and pathway enrichment in *Ldb1(−/−)* embryonic stem cells and embryoid bodies

To investigate the role of LDB1 in regulating proliferation and differentiation, we conducted transcriptome analysis for WT and *Ldb1(−/−)* ESC, EB and differentiated erythroblasts by RNA-seq. There were distinct patterns of gene expression changes between WT and *Ldb1(−/−)* cells at each of the three stages, as evident from hierarchical clustering and principal component analysis (PCA) (Fig. S3A, B). Volcano plots indicated 1898 dysregulated genes (515 downregulated, 1,383 upregulated) in *Ldb1(−/−)* ESC compared to controls, and 829 dysregulated genes (322 downregulated, 507 upregulated) at the EB stage (*P*adj <0.05, FC>1.5) (Fig. 3A, B). Most expressed genes were shared by WT and *Ldb1(−/−)* cells at ESC and EB stages, with only a small percentage expressed only in WT or *Ldb1(−/−)* (Fig S3C, D). RNA-seq results confirmed the reduced transcription of *Oct4* and *Sox2* we had observed by RT-qPCR for *Ldb1(−/−)* ESC but not for Nanog, suggesting that, in this case, the effect of LDB1 loss may be post-transcriptional (Fig. S3E). DEGs that were shared upon LDB1 loss in ESC and EB were dysregulated similarly, i.e. up or down regulated in both (Fig. S3F)

**Figure. 3.**
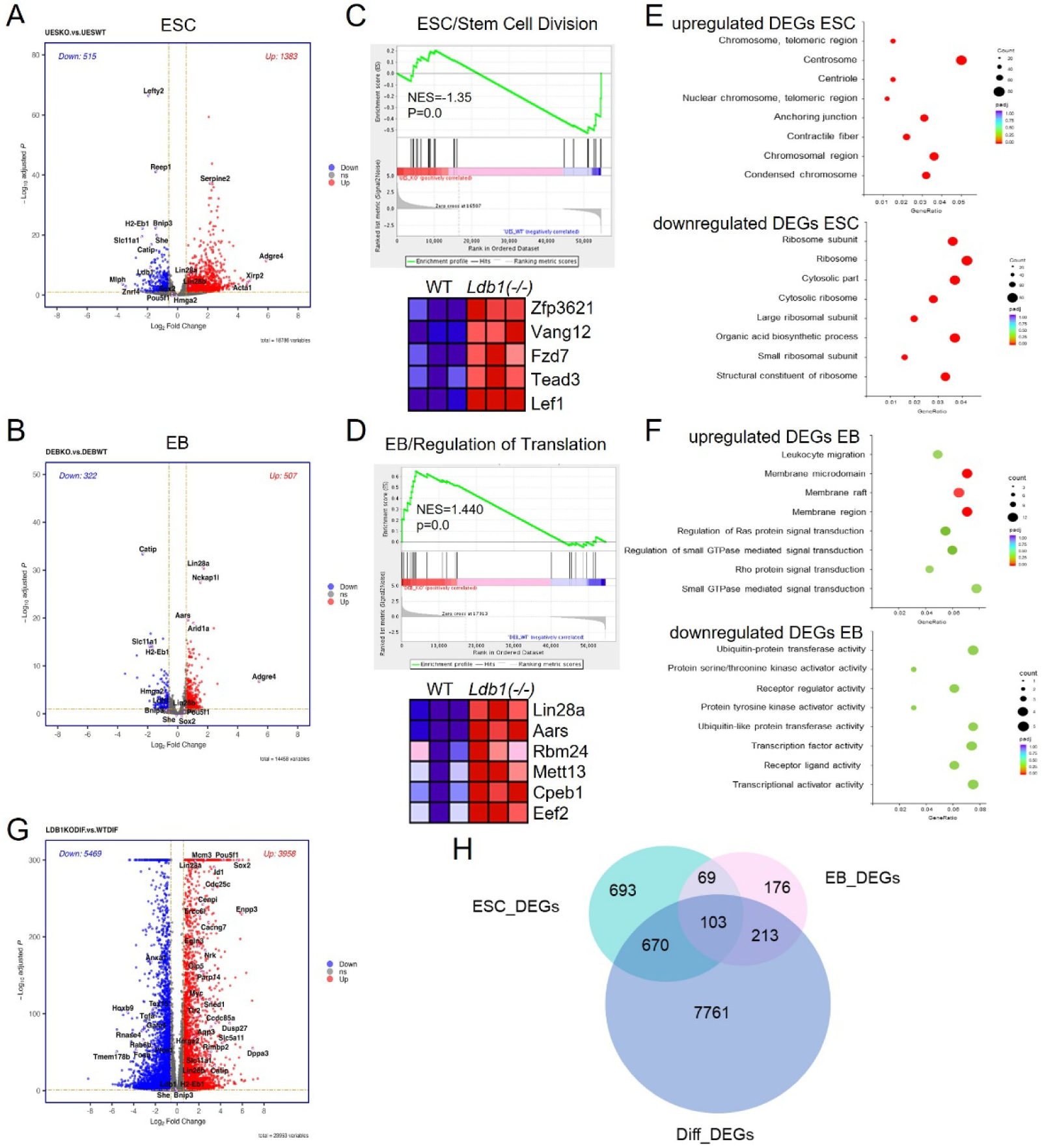
Altered **transcriptome after loss of LDB1 in ESC and EB.** (A, B) Volcano plots show log2-fold change (X-axis) and statistical significance (Y-axis) of gene expression changes between WT and *Ldb1(−/−)* ESC and EB (*P*adj <0.05), FC>1.5. N=2. (C, D) GSEA enrichment of specific biological processes in WT or *Ldb1(−/−)* ESC and EB. Heatmaps (right) highlight specific genes associated with stem cell division or cytoplasmic translation differentially enriched in *Ldb1(−/−)* ESC or EB. (E, F) Bubble plots of GO molecular functions enriched in up-or downregulated DEGs from (A) and (B) in ESC and EB. (G) Volcano plot of gene expression changes between WT and *Ldb1(−/−)* ESC and EB (*P*adj <0.05). N=2. (H) Venn diagram comparing DEGs from ESC, EB and erythroblast stages after loss of LDB1. *P*adj<0.05, FC>1.5.

To assess the potential physiological relevance of dysregulated genes in *Ldb1(−/−)* cells, Gene Set Enrichment Analysis (GSEA) was carried out, revealing enrichment of somatic stem cell division genes in *Ldb1(−/−)* ESC (FDR <0.05) (Fig. 3C). The accompanying heatmap displays expression changes for representative genes, including WNT pathway *Lef1* and *Fzd7*, suggesting potential effects on the stem cell phenotype due to altered expression of key signaling molecules. In EB cells, LDB1 loss was associated with enrichment of proteins involved in cytoplasmic translation, including LIN28 (Fig. 3D). Gene ontology (GO) analysis of DEGs after LDB1 loss revealed an enrichment of genes with molecular functions related to chromosome condensation among upregulated DEGs in ESC and of ribosome-related functions among downregulated DEGs (Fig. 3E). In EB, upregulated and downregulated DEGs upon LDB1 loss were enriched for membrane function or protein processing, respectively, but the enrichments were of low significance (Figure 3F).

After further differentiation of WT or *Ldb1(−/−)* ESC to erythroblasts, gene dysregulation was the most dramatic with 5,469 up– and 3,958 downregulated genes (*P*adj<0.05, FC>1.5) (Fig. 3G). A very large fraction of DEGs (85%) corresponded to erythroid fingerprint genes (50). Most DEGs at this stage (89%) were unique to the differentiated cells and not shared by ES or EB (Fig. 3H). These results are consistent with the large effect of LBD1 loss on erythroid cell differentiation markers seen in Fig. S2C, D and with the known role of LDB1 enhancers in erythroid cell differentiation (51,52). Overall, this analysis highlights significant changes in expression of pluripotency factors in *Ldb1(−/−)* ESC and of dysregulation of NODAL (*Lefty2*), WNT (*Lef1*) and *LIN28* pathway genes that may contribute to the effects of LDB1 loss on proliferation and differentiation of ESC.

### Disruption of *Ldb1* leads to dysregulation of *Lin28b*-mediated stem cell self-renewal activity

Previous studies indicated the crucial involvement of LIN28 in stem cell self-renewal via the *let-7* microRNA pathway and mRNA translation regulation, and LIN28a and LIN28b are also key factors implicated in organismal growth, metabolism, and tissue development (42). To validate the expression changes of *Lin28* in *Ldb1(−/−)* ESC, we performed RT-qPCR and western blot analyses. *Ldb1(−/−)* ESC exhibited a pronounced increase in *Lin28a* and *Lin28b* expression (Fig. 4A, Fig.S4A). Western blotting and immunofluorescent staining of *Ldb1(−/−)* ESC indicated a significant increase in LIN28b protein compared to wild-type ESC (Fig. 4B, C).

**Figure. 4.**
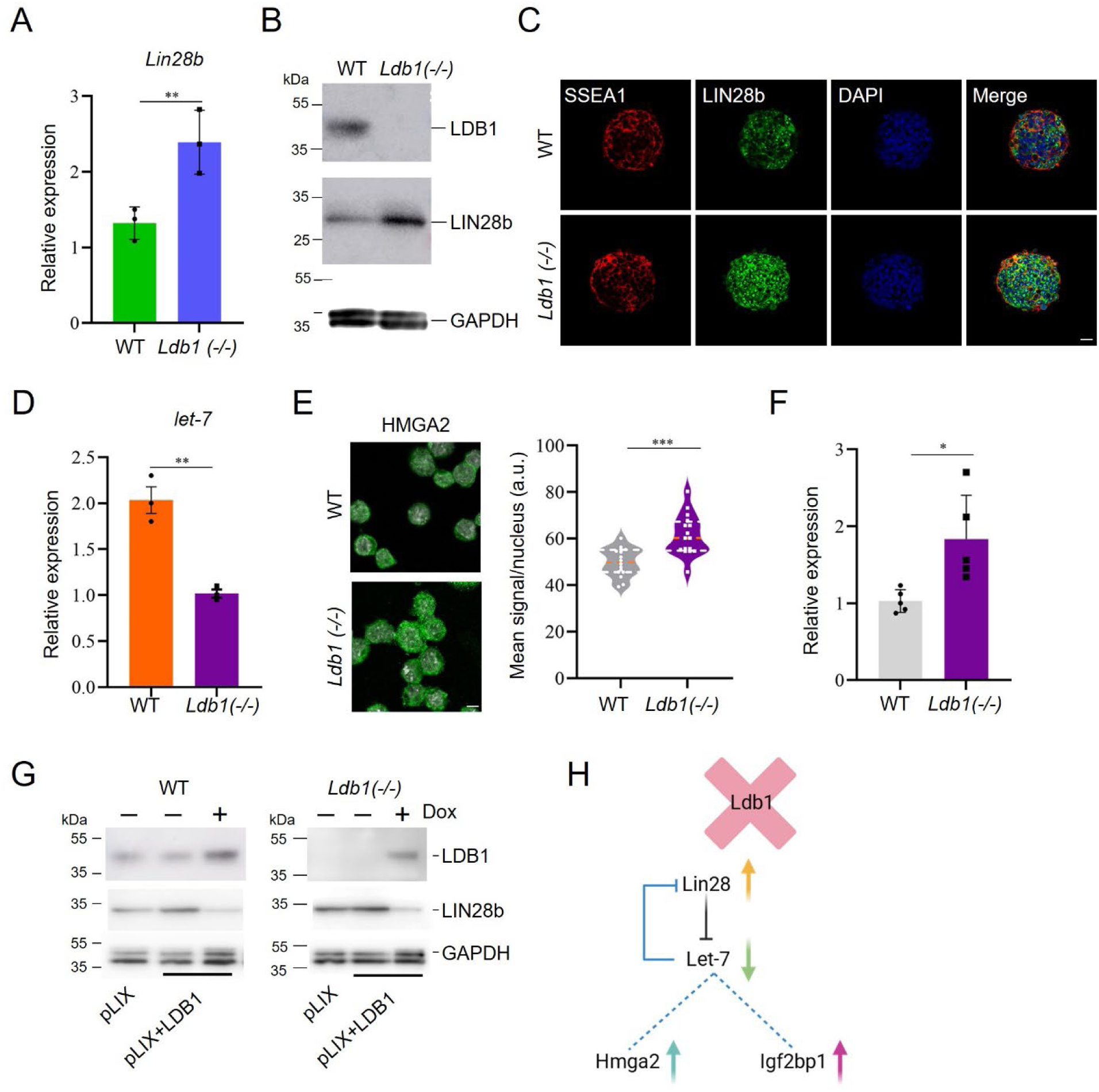
LIN28 is **upregulated and the Let7 pathway is dysregulated in *Ldb1(−/−)* ESC.** (A) RT-qPCR analysis of *Lin28b* mRNA in WT and *Ldb1(−/−)* ESC. (B) Western blot analysis of LIN28b in WT and *Ldb1(−/−)* ESC. (C) Representative immunofluorescent images of LIN28b in WT and *Ldb1(−/−)* ESC. Scale bar, 20 μm. (D) RT-qPCR of *mir-Let7a* in WT and *Ldb1(−/−)* ESC. (E) Representative immunofluorescent images of HMGA2 in WT and *Ldb1(−/−)* ESC. Scale bar, 5 μm. Quantitation of HMGA2 protein immunofluorescence intensity using image J is shown on the right. (F) RT-qPCR analysis *Hmga2* expression in WT and *Ldb1(−/−)*. (G) Western blot analysis showing the recovery of WT LIN28b protein levels in *Ldb1(−/−)* ESC upon LDB1 overexpression. (H) Diagram depicts changes in LIN28/Let-7 and target proteins after loss of LDB1. Data quantitation represents mean ± SEM, n=3 biological replicates. Statistical significance assessed using two-tailed Student’s t-test. * p<0.05, ** p<0.01, *** p<0.001.

LIN28 regulates the biogenesis of *let-7* microRNA (53). Notably, *let-7a* expression *7*, displayed increased protein and both mRNA and nascent RNA expression in *Ldb1(−/−)* ESC (Fig. 4E, F, Fig. S4B, C). We assessed the expression of insulin-like growth factor 2 mRNA binding proteins (*Igf2bp*), also *let-7* targets and known to be regulated by HMGA2 (46). Although we examined *Igf2bp1*, *Igf2bp2*, and *Igf2bp3*, only *Igf2bp1* expression decreased in *Ldb1(−/−)* ESC (Fig. 4D). *Hmga2,* which is targeted for degradation by *let-*was increased in the *Ldb1(−/−)* ESC (Fig. S4D).

To validate the regulatory role of LDB1 in LIN28b expression in ESC, we employed a Tet-inducible LDB1 expression system (Fig. S4E, F). Following transient transfection of the Tet-inducible *Ldb1* vector into WT and *Ldb1(−/−)* ESC, incubation was continued for 48 hours in the presence of doxycycline. The increase in LIN28b observed in *Ldb1(−/−)* cells was reversed upon LDB1 overexpression to levels similar to WT cells, providing supportive evidence of LDB1-mediated regulation of LIN28b (Fig. 4G). The results suggest that the effects of LDB1 loss in ESC and EB may, at least in part, result from the upregulation of *Lin28* and the subsequent effect on the LIN28-mediated *Let-7* signaling pathway (Fig. 4H).

### Global loss of chromatin accessibility in *Ldb1(−/−)* ESC and EB

Previous studies have highlighted global remodeling of the epigenomic state during transcription factor-induced cell developmental stage transitions (1,5,54). To assess the differential accessibility of chromatin upon loss of LDB1, we conducted a genome wide analysis using Assay for Transposase-Accessible Chromatin sequencing (ATAC-seq) in WT and *Ldb1(−/−)* ESC and EB (FDR<0.05, FC>2). Heatmaps show that genome wide, ATAC-seq peaks were dramatically reduced in ESC by loss of LDB1 and somewhat less so in EB (Fig.5A). This trend was magnified when promoter or enhancer localized ATAC-seq peaks were clustered (Fig. S5A). Globally, there was a loss of 39,624 accessible peaks and 97,849 accessible peaks comparing WT to *Ldb1(−/−)* ESC or EB, respectively (Fig. S5B). The DiffBind software identified 5,173 differentially accessible sites after LDB1 loss, all but 10 of which lost accessibility in *Ldb1(−/−)* ESC and 8,856 differentially accessible peaks after LDB1 loss in EB, 84% of which decreased (Fig. 5B).

**Figure. 5.**
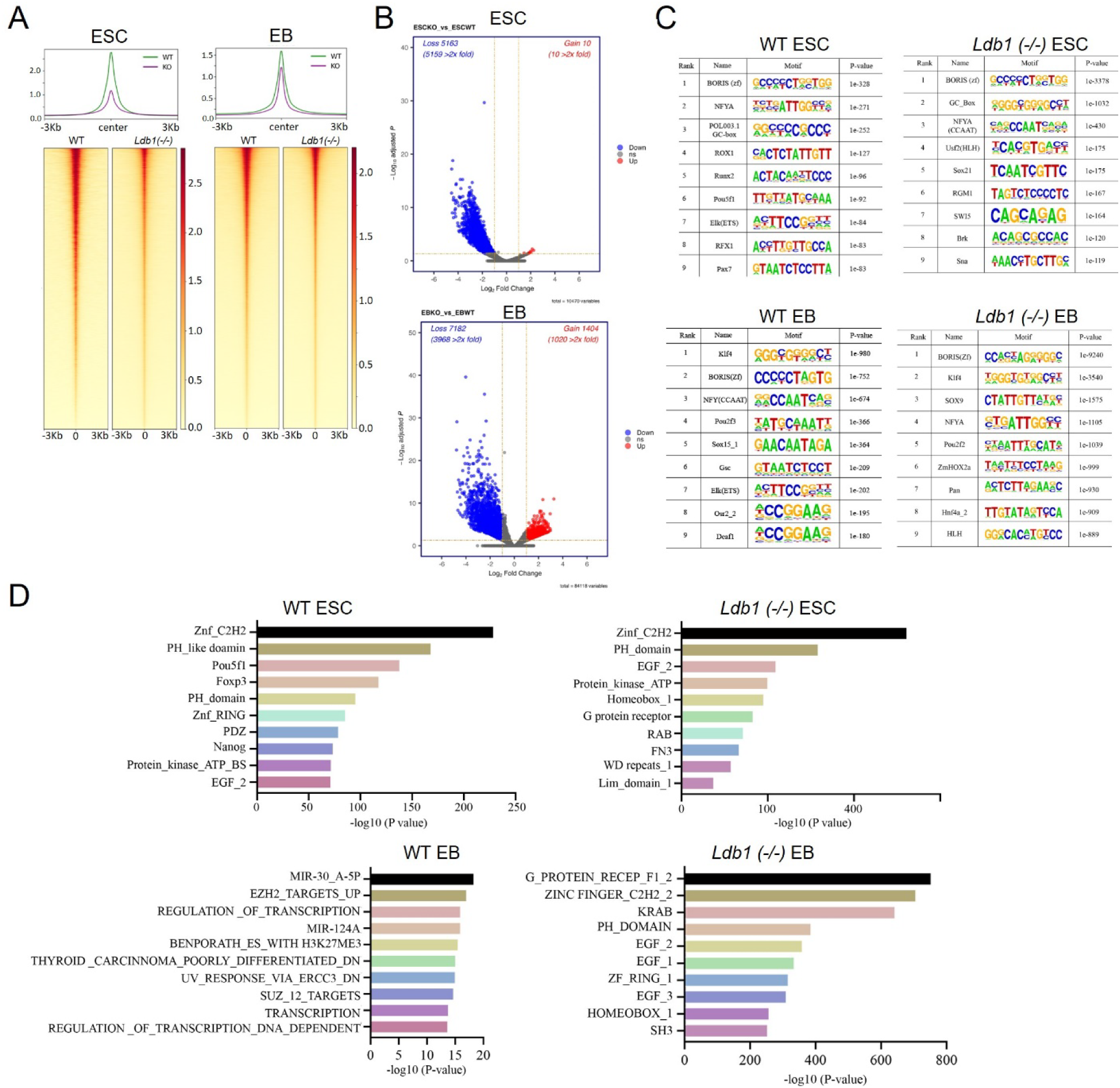
ATAC-seq r**eveals global decrease in chromatin accessibility *Ldb1(−/−)* ESC and EB compared to WT cells.** (A) Heatmaps show chromatin accessibility genome wide in WT and *Ldb1(−/−)* ESC and EB using ATAC-seq. N=2 biological replicates. (B) DiffBind analysis of chromatin accessibility for *Ldb1(−/−)* ESC and EB compared to WT. Red dots indicate significant change in accessibility. (C) Motif analysis identifies the transcription factor binding sites enriched in the accessible chromatin regions of WT and *Ldb1(−/−)* and ESC and EB. (D) Bar graphs showing the enriched protein families associated with the identified accessible regions in WT and *Ldb1(−/−)* ESC and EB.

An analysis using MEME-ChIP of motifs enriched in accessible peaks of ESC and EB WT and *Ldb1(−/−)* cells prominently revealed BORIS, the stem cell CTCF homologue, at both developmental stages and regardless of LDB1 status (Fig. 5C). Interestingly, the *Pou5f1* (*Oct4*) motif was strongly enriched at accessible chromatin regions in WT ESC but not in *Ldb1(−/−)* cells, suggesting that LDB1 loss influences sites of OCT4 occupancy. By contrast, the *Klf4* motif was strongly enriched at accessible regions in WT EB and was unaffected after LDB1 loss (Fig. 5C). Pou5f1 (OCT4) and Nanog protein family members were among those enriched at accessible chromatin regions in ECS, while several microRNA families and transcription related protein families were enriched at these regions in EB (Fig. 5D). After *Ldb1* deletion, these enrichments were lost. These data show that the loss of LDB1 results in a significant decrease in chromatin accessibility globally at enhancers and may have a broad effect on pluripotency gene occupancy in ESC and EB.

### Role of LDB1 in enhancer function during development

The data so far suggest a possible relationship between pluripotency factors and LDB1 chromatin occupancy. To determine potential chromatin targets of LDB1, we performed LDB1 ChIPmentation in WT ESC. MACS2 and IDR filtering (*P*val<0.05) called 1048 high confidence LDB1 peaks of which about 11% were at promoters and 84% were intergenic or intronic, consistent with the known enrichment of LDB1 at enhancers (51,52) (Fig. S6A). MEME-ChIP motif analysis showed enrichment at LDB1 peaks for the pluripotency protein KLF4 motif, for SOX family members and for TEAD family members, which are enhancer proteins in stem cells (Fig. 6A). Strikingly, LDB1 localization in ESC overlapped with that of pluripotency factors from published data at 504 sites in ESC, which represent 85% of all the reported OSN overlapped sites (Figure 6B). Moreover, LDB1 was enriched at ESC super enhancers (SE) in these cells and enrichment was stronger than at typical enhancers (TE) (Fig.6C). LDB1 occupancy overlapped with 30% of the reported 231 SEs in ESC (33).

**Figure. 6.**
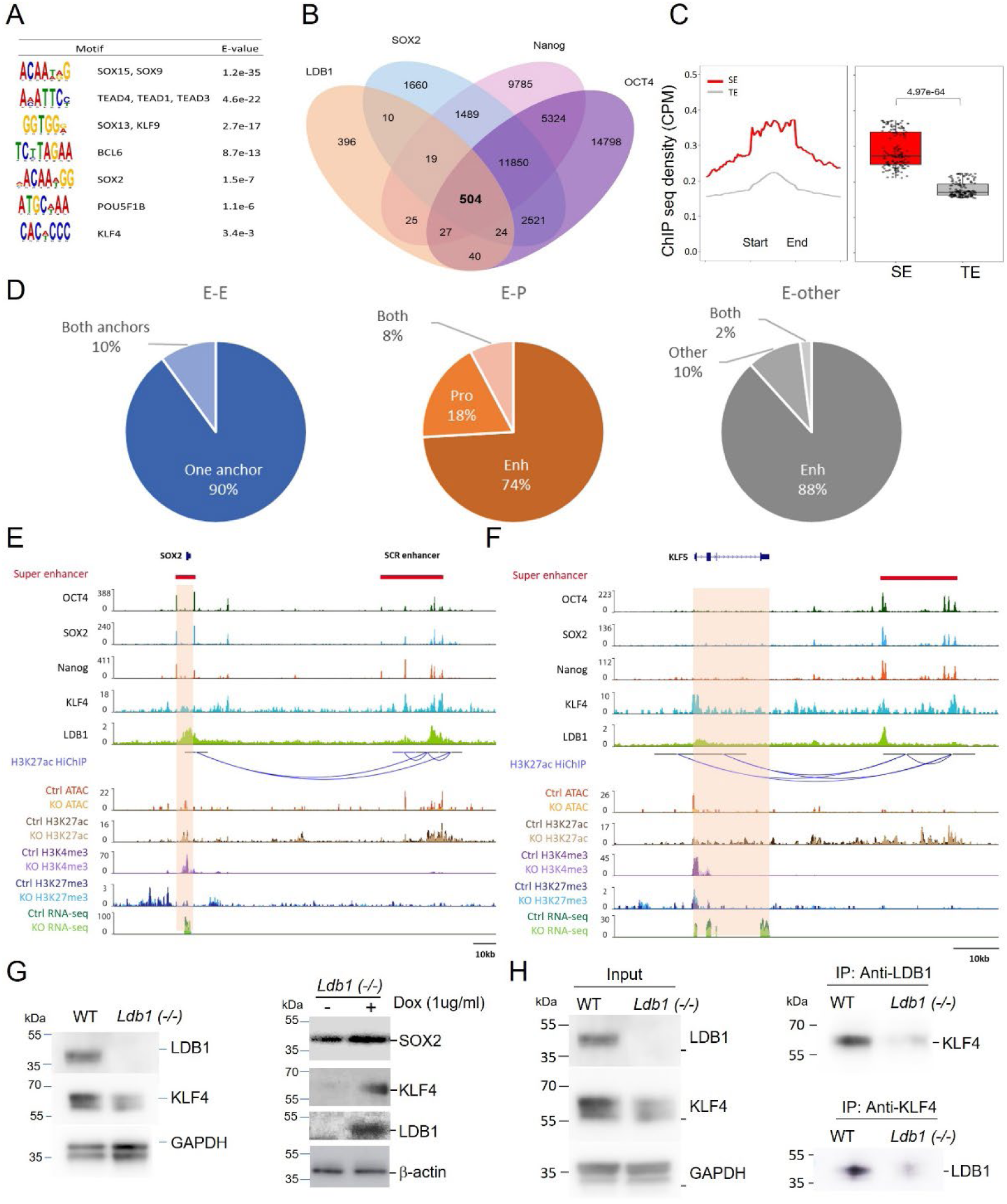
Co-occupancy of **LDB1 and OSN at super enhancers in ESC and EB.** Motifs enriched at LDB1 peaks in ESC. (B) Venn diagram of overlap of LDB1 peaks with OCT4, SOX2, and Nanog in ESC. (C) Enrichment of LDB1 peaks in super enhancers (SE) and typical enhancers (TE) from Whyte et al. (D) Pie charts of LDB1 occupancy at loop anchors from Di Giammartino in ESC (39). (E) Genome browser view of the *Sox2* locus. OCT4, SOX2, Nanog and KLF4 are from (34) and H3K27ac HiChIP data are from (39). (F) Genome browser view of the *KLF5* locus with tracks as in panel E. (G) Left, Western blot analysis of LDB1 and KLF4 in WT and *Ldb1(−/−)* ESC. Right, after LDB1 expression by doxycycline inducible vector in *Ldb1(−/−)* ESC. (H) Immunoprecipitation with antibodies to LDB1 or KLF4 and western blotting with the reciprocal antibody.

Enhancers establish proximity to target genes for activation. To determine long range interactions engaged in by LDB1, we integrated our LDB1 ChIPmentation data with enhancer looping data derived from H3K27ac Hi-ChIP in ESC (39). There were 7,285 loops that involved at least one LDB1-occupied loop anchor. Analyzing these results together with enhancers called in ESC from Whyte *et al.* (34), we separated the loops into those involving two enhancers (E-E), an enhancer and a promoter (E-P), or an enhancer engaged in a loop with a fragment that was not defined as a promoter or an enhancer (E-other). In all groups, most loops had LDB1 at only one anchor with 10% or less having LDB1 at both anchors (Fig. 6D). For E-P loops, 82% had LDB1 at the enhancer anchor and 26% at the promoter anchor. These results are consistent with results of LDB1 long-range enhancer interactions in erythroid cells (52).

The *Sox2* locus is an illustrative example of co-localization of LDB1, OCT4, SOX2 and KLF4 both at *Sox2* and at the SCR enhancer (Fig. 6E)(55). H3K27ac HiChIP shows the interaction between the gene and enhancer (39). Chromatin accessibility and H3K27ac, determined by Cut & Tag, were reduced across the gene and enhancer when the gene KLF4 has been implicated in pluripotency gene 3D enhancer long range interactions (37,39). Our data show that not only is LDB1 a positive regulator of KLF4 but it is prominently co-localized with KLF4 at OSN-occupied ESC enhancers (28%). Earlier work found that in erythroid cells LDB1 was often co-localized with the related Kruppel-like factor KLF1, although typically LDB1 cooperates with LIM-homeodomain and LIM only proteins to occupy chromatin and loop enhancers (13,51,56). To further explore the relationship between KLF4 and LDB1 in ESC, we observed by western blot the reduction in precipitated KLF4 protein upon LDB1 loss and rescue upon LDB1 overexpression (Fig, 6G). We then used immunoprecipitation and western blotting to provide evidence that LDB1 and KLF4 can interact in ESC, whereas no interaction was observed between LDB1 and OCT4 (not shown) (Fig. 6H). Together, these results support that LDB1 functions together with pluripotency factors OCT4, SOX2, Nanog and KLF4 at SE and TE to regulate a subset of genes in ESC and in EB, including the pluripotency factors themselves.

### Deletion of *Ldb1* reduces hematopoietic stem cells in bone marrow

Our experiments differentiating *Ldb1(−/−)* ESC to erythroblasts indicated loss of LDB1 impeded acquisition of erythroid cell surface markers and β-globin transcription. The role of LDB1 in differentiation in an *in vivo* context remained unexplored. To investigate further, we bred *Ldb1* conditional knockout mice using an interferon-inducible *Mx1-Cre* allele and *Ldb1^flox/flox^* mice (57) (Fig. S7A-C). We treated *Mx-Cre:Ldb1^flox/flox^* mice and controls with polycytidylic acid (pIpC) at three time points before isolating bone marrow and peripheral blood for examination of erythroid maturation in the presence or absence of LDB1 (Fig. 7A). We conducted immunofluorescent staining and FACS analysis for representative marker CD15 for peripheral blood mononuclear cells (PBMC). The results revealed an approximately 3-fold reduction in CD15 in PBMC of *Ldb1* conditional null mice (Fig. 7B).

**Figure. 7.**
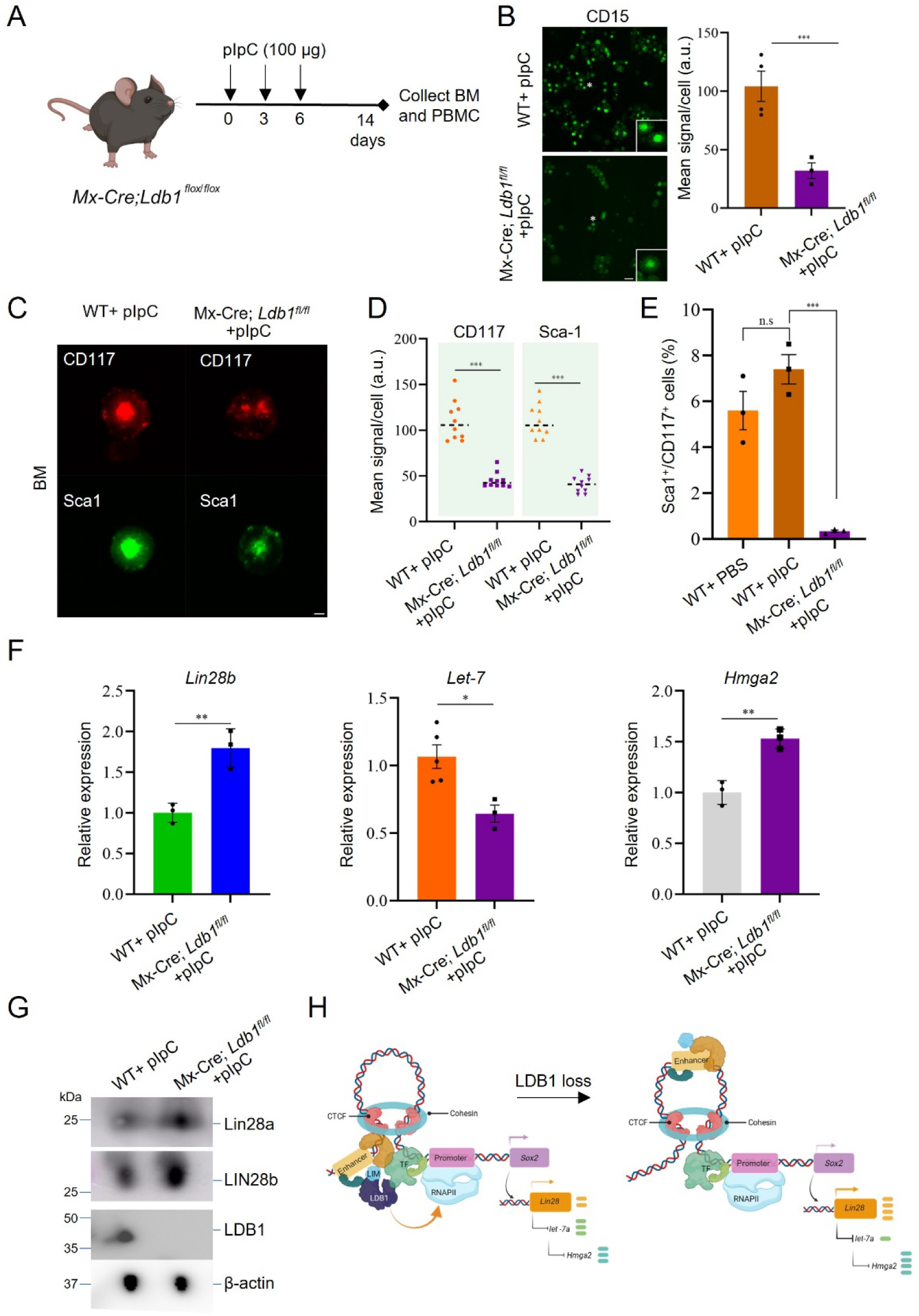
Conditional loss of LDB1 in mouse bone marrow of *Mx-Cre; Ldb1^flox/flox^* mice recapitulates the effect of LDB1 loss on LIN28 in ESC. (A) pIpC injection protocol and bone marrow isolation from *Mx-Cre; Ldb1^flox/flox^* Mice. (B) Immunofluorescent staining for CD15 in PBMCs from WT+ pIpC and *Mx-Cre; Ldb1^flox/flox^* +pIpC Mice. Cell count: 20 cells/slide. N=3 mice. Scale bar, 5 µm. Right, quantitation of CD15 expression via image J. (C) Immunofluorescent staining of CD117 and Sca-1 in BM from WT+ pIpC and *Mx-Cre; Ldb1^flox/flox^* +pIpC Mice. Cell count: 10 cells/slide. N=3 mice. Scale bar, 5 µm. (D) Quantitation of CD117 and Sca-1 mean signal per cell using image J for FACS sorted cells shown in Fig. S7D from WT+ pIpC and *Mx-Cre; Ldb1^flox/flox^* +pIpC Mice. (E) FACS analysis of BM using Sca-1 and CD117 markers displaying the percent of double positive cells from WT+ pIpC and *Mx-Cre; Ldb1^flox/flox^* +pIpC Mice. (F) RT-qPCR analysis of *Lin28b*, *miR-Let7* and *Hmga* expression in BM from WT+ pIpC and *Mx-Cre; Ldb1^flox/flox^* +pIpC Mice. (G) Western blot analysis of LIN28a and b in liver of WT+ pIpC and *Mx-Cre; Ldb1^flox/flox^* +pIpC mice. (H) Model of the effect of Ldb1 loss in ESC is depicted. Data quantitation represents mean ± SEM, n=3 biological replicates. Statistical significance assessed using two-tailed Student’s t-test. * p<0.05, ** p<0.01, *** p<0.001.

Next, we performed immunofluorescent staining and FACS analysis on isolated bone marrow cells (Fig. 7C, Fig. S7D). Markers associated with hematopoietic progenitors (Sca-1^+^ and CD117^+^) that are crucial for bone marrow function exhibited notable decreases in mean signal per cell and number of double positive cells in *Mx1-Cre;* null mice (Fig. 7D, E). These observations support an impact of LDB1 on erythroid differentiation during development. Finally, we performed RT-PCR for *Lin28b* and downstream targets *Let7a* and *Hmga2* using isolated bone marrow cells from *Mx1-Cre; Ldb1^flox/flox^* mice. We observed elevated *Lin28b* transcription, reduced *Let-7a* transcription and elevated *Hmga2* transcription, aligning with our prior results in ESC (Fig. 7F). Furthermore, LIN28a/b proteins were elevated in bone marrow cells after loss of LDB1 in *Mx1-Cre; Ldb1^flox/flox^* mice (Fig. 7G). Collectively, these findings emphasize the influence of LDB1 on LIN28/*Let-7*/HMGA2 signaling in the context of erythropoiesis *in vivo*.

## DISCUSSION

In this study, we investigated the role of LDB1 in ESC and its impact on aspects of stem cell biology and development to EB and erythroblasts. We found that the loss of LDB1 resulted in significantly increased proliferation and stem cell activity observed in *Ldb1(−/−)* ESC. EB formation from *Ldb1(−/−)* ESC was impaired and differentiation of disrupted EBs to erythroblasts was delayed. The results underscore an essential role for LDB1 in stem cell maintenance and their ability to engage in lineage choice. The regulation of pluripotency factors is crucial for stem cell self-renewal and to ensure their ability to differentiate into various cell lineages during development (58). Loss of LDB1 resulted in changes in the abundance of key transcription factors, including SOX2, OCT4, and KLF4 but, more broadly, NODAL, WNT and LIN28 pathway genes were affected by LDB1 loss. Underlying these effects was substantial co-occupancy of LDB1 with pluripotency factors at ESC super enhancers and effects of LDB1 loss on chromatin accessibility. Thus, LDB1 has a previously unappreciated role in maintaining pluripotency and influencing the fate of ESC.

OCT4, SOX2 and Nanog are known to positively regulate *Lin28* and *Let-7g* microRNA primary transcripts by occupying their promoters (44,59). Mechanistically, mature *Let-7* contributes to degradation of LIN28, while LIN28 blocks the maturation of *Let-7g* in a feedback regulatory loop. We found that LDB1 positively regulates OCT4, SOX2 and Nanog, which are reduced upon LDB1 loss. This change contributes directly, and probably indirectly, to an imbalance of LIN28 and mature *Let-7*, in which *Let-7* is decreased and LIN28 is elevated. LIN28 is known to promote pluripotency. Based on these results, we propose that LDB1 is important for the interplay of miRNAs and pluripotency factors in ESC to both maintain stemness and the differentiated state.

We speculate that after the EB stage there is a shift from LDB1 enhancer cooperation with pluripotency factors at enhancers to function with erythroid factors GATA1, TAL1 and LMO2. In this scenario, KLF4, which shares many sites of occupancy with OCT4, SOX2 and Nanog, and is critical for enhancer looping in ESC, may function in mediating LDB1 association with chromatin in ESC and EB, analogous to a proposed function of *Kruppel* family member KLF1 in erythroid cells (39,51). In erythroid cells and other cell types, LIM-HD proteins and LIM only proteins have been shown to provide this function (13). In ESC it remains to be investigated whether there is a LIM domain protein that functions with LDB1 in enhancer looping.

In the model (Fig. 7H), we speculate that LDB1, interacting with KLF4 and/or an as yet unknown protein complex including a LIM domain protein, binds at targets, for example, the *Sox2* promoter and enhancer, contributing to long range interaction between them to increase transcription, beyond the contribution of cohesin and mediator (32). LDB1 has both cohesin-related and cohesin independent enhancer looping activity (60).

Dimerization of LDB1 may initiate or stabilize chromatin loop formation to regulate *Sox2* (21,61). The absence of LDB1 decreases SOX2, disrupts this mechanism, and upregulates the promoter of pluripotency LIN28 through the mechanism of Let-7/LIN28 linkage (44). LDB1-mediated enhancer function, thus, critically balances the equilibrium between *Lin28* expression to promote pluripotency and cellular differentiation in ESC. The influence of LDB1 in ESC may extend farther to regulatory networks that involve, in addition, NODAL and WNT signaling components (41).

We further investigated the impact of LDB1 deficiency on erythroid differentiation. Our results revealed impaired erythroblast formation potential in *Ldb1(−/−)* ESC, as indicated by reduced expression of lineage-specific cell surface markers CD71 and Ter119 and the delayed formation of erythroblasts. Vastly more DEGS were observed in LDB1*(−/−)* differentiated erythroblasts than in ESC or EB and 85% of these DEGs were erythroid fingerprint genes (50). This result is consistent with the substantial role LDB1 enhancers play in erythroid cells (51,52). Furthermore, our *in vivo* experiments show that LDB1 is indispensable for proper erythroid differentiation in bone marrow in adults. Yu, *et al*. (27) had documented, using the proximity ligation assay, that GATA1/LDB1 complexes become increasingly important in terminal erythroid differentiation. Other work in which GATA1 HiChIP was performed, showed that cell-type specific gene expression becomes increasingly dependent on LDB1 enhancers with developmental stage from embryonic to adult erythropoiesis (26). Our work specifically suggests that the targets of LDB1 enhancer looping complexes in erythroid cells are different from the subset of enhancer-gene pairs that are LDB1-dependent in ESC and EB.

Our genome-wide analysis of chromatin accessibility revealed the impact of LDB1 deficiency on chromatin remodeling (33). The significant reduction in chromatin accessibility, particularly at enhancers, indicates that enhancers are largely reprogrammed in *Ldb1(−/−)* ESC and EB. The stem cell form of CTCF, BORIS, was enriched at accessible chromatin peaks in ES and EB regardless of the presence of LDB1, suggesting they may cooperate, for example, at TAD borders where CTCF is enriched. LDB1 has known interactions with CTCF (52). However, the motifs for OCT4 and KLF4 were enriched at accessible chromatin peaks only in WT ESC and EB and the enrichment was lost when accessibility globally decreased in *Ldb1(−/−)* cells. Since LDB1 co-occupies many sites in chromatin together with these factors, this suggests that LDB1 may influence the occupancy of OCT4 and KLF4 in ESC and EB.

In conclusion, our findings provide insight into the molecular mechanisms by which LDB1 influences chromatin structure, gene expression, and critical aspects of stem cell biology. Further investigations are warranted to elucidate the precise molecular pathways and downstream targets involved in LDB1-mediated regulation of stem cell activity and gene expression. Integration of the LDB1 regulatory network into the broader context of stem cell biology promises to advance our knowledge of stem cell fate decisions and inform the potential application of stem cells in treatment of genetic blood diseases and cancers.

## DATA AVAILABILITY

ChIP-seq data was downloaded from the GEO website using the following accession numbers: GSE44286 (OCT4, SOX2 and Nanog ChIP-Seq) (Whyte et al. 2013), GSM288354 (KLF4 ChIP-Seq) (Whyte et al. 2013), and GSE113339 (H3K27ac Hi-ChIP) (Di Giammartino et al. 2019).

All sequence data acquired in the work has been submitted to GEO under the following accession numbers: GSE282372 (RNA-seq, reviewer token, uparkowgpdgjvgx) GSE282373 (ChIPmentation, reviewer token, wdgzwaoqpbeztyj) GSE282374 (Cut-&-Tag, reviewer token, qjevgoucbjevpsv) GSE282375 (ATAC-seq, reviewer token, ajslqkgizlgdvwj)

## SUPPLEMENTARY DATA

Supplementary Figures S1-S7; Supplementary Table S1. Primers used in this study. Table S2. Antibodies used in this study.

Supplementary data are available at NAR Online.

## Supporting information

LDB1 manuscript

## ACKNOWLEDGEMENTS

We thank Dr. Pedro Rocha for helpful comments on the manuscript. We acknowledge the NIDDK Genome Core and NHLBI Sequencing and Genomics Core Facility for sequencing. This work utilized the computational resources of the NIH HPC Biowulf cluster (https://hpc.nih.gov). We thank Dr. Yangu Zhou for *Mx-Cre:Ldb1^flox/flox^* mice.

## DECLARATION OF INTERESTS

The authors declare that they have no competing interests

## FUNDING

This work was funded by the Intramural program of the National Institute of Diabetes and Digestive and Kidney Diseases, NIH (DK 075033 to A.D.)

## Notes

### Competing Interest Statement

The authors have declared no competing interest.

