## Supplementary material for "LDB1 regulates gene expression and chromatin structure in pluripotency and lineage differentiation": LDB1 manuscript

#### SUPPLEMENTARY INFORMATION

##### Supplementary Methods

##### Supplementary Figures

Figure S1-S7.

##### Supplementary Tables

Table S1. Primers used in this study. Table S2. Antibodies used in this study.

**Supplementary Methods**

*RNA-seq data analysis*

RNA-seq fastq files were trimmed using the cutadapt program (v2.7) with the settings "-a AGATCGGAAGAGCACACGTCTGAACTCCAGTCA -A AGATCGGAAGAGCGTCGTGTAGGGAAAGAGTGT -q 20 -m 25". Sequencing library quality was assessed with fastqc (v0.11.8) (https://qubeshub.org/resources/fastqc) with default parameters. Preprocessed reads were then aligned to the mouse reference genome GRCm38 from Gencode M18 using hisat2 (v2.1.0) [1] with default parameters. Aligned reads were then mapped to gene features using subread featureCounts (v1.6.4) [2] with parameters "-p -s 0 -T 1". Differential expression between groups of samples was tested using R (version 3.5.1) with DESeq2 package (v1.22.1) [3]. Transcript quantitation was performed with salmon (v0.14.2) [4] with parameters "--libType=A --gcBias --seqBias --validateMappings". QC data were summarized with multiQC [5]. Coverage tracks were generated using the deepTools (v3.3.1) bamCoverage with "--minMappingQuality 20 --smoothLength 10 --normalizeUsing BPM".

*Processing and analysis of ChiPmentation*

Single-end raw fastq files were trimmed using the cutadapt program (v2.7) with the settings "-a CTGTCTCTTATA -q 20 --minimum-length 25". Preprocessed reads were aligned to GRCm38 genome using Bowtie2 (v2.3.5) with the "--no-unal" option. After removing non-uniquely mapped reads using samtools (v1.9) with "-q 20" [6] and duplicated reads using picard MarkDuplicates (v2.21.4) with "REMOVE_DUPLICATES=true", final reads were retained for peak calling using MACS2 [7,8] (v2.2.7.1) with the settings "-f BAM --keep-dup all -g 2.3e9" with the input sample as control at default cutoffs. Browser track files were generated using the deepTools [9] (v3.5.2) bamCoverage with "--minMappingQuality 20 --ignoreDuplicates --normalizeUsing CPM --extendReads 300".

In downstream analyses, reads mappable to mm10 blacklist regions [10] or non-canonical chromosomes were further removed from peak files using BEDTools [11]. Irreproducible discovery rate (IDR) software (v2.0.3) [12] was deployed to measures consistency between replicates with p.value 0.05 as cutoff. Motif analysis on the selected reproducible subset of peak sequences centered around summits was performed using the MEME-ChIP software [13] with the databases “uniprobe_mouse.meme” and “JASPAR2018_CORE_vertebrates_non-redundant.meme” and parameters “-ccut 100 -order 3 -meme-mod zoops -meme-minw 6 -meme-maxw 30 -meme-nmotifs 5 -meme-searchsize 100000 -dreme-e 0.05 -centrimo-score 5.0 -centrimo-ethresh 10.0”.

*ATAC-seq data analysis*

Paired-end raw fastq files were preprocessed using the cutadapt program (v4.4) with the settings “-a CTGTCTCTTATACACATCT -A CTGTCTCTTATACACATCT --nextseq-trim 20 --overlap 6 --minimum-length 25” to trim adapter and low-quality sequences. Preprocessed reads were aligned to the mouse genome GRCm38/mm10 available at Gencode M18 [14] using Bowtie2 (v2.5.1) [15] with the settings “-X 2000 --no-unal”. Uniquely mapped, properly paired, non-supplementary, non-mitochondrial alignments were retained using samtools [6] view command with options “-F 2048 -f 2 -q 20” piped with “grep -v chrM”. Duplicated reads were cleaned using picard MarkDuplicates (v2.27.5) (http://broadinstitute.github.io/picard) with “REMOVE_DUPLICATES=true”. Peaks were called using MACS2 [7,8] (v2.2.7.1) with the settings “-f BAMPE --keep-dup all -g 2.3e9” at default cutoffs. Browser track files were generated using the deepTools [9] (v3.5.2) bamCoverage with “--minMappingQuality 20 --ignoreDuplicates --normalizeUsing CPM --extendReads 300”. Identification of differential accessibility regions between two biological conditions was performed with R package “DiffBind” (v3.8.4). In downstream annotation and visualization analyses, reads mappable to mm10 problematic genome regions defined by the ENCODE Blacklist [10], or aligned on non-canonical chromosomes, were further removed from BAM files using bedtools intersect with the “-v” option.

The filtered bam files of biological replicates were merged per condition using the samtools merge function. Normalized bigwig signals per condition were calculated with bamCoverage “--binSize 50 --normalizeUsing CPM --extendReads –ignoreDuplicates”.

Signal ratio (log2) between KO and WT was calculated via bamCompare with parameters "--operation log2 --scaleFactorsMethod None --normalizeUsing CPM --extendReads --ignoreDuplicates --binSize 50". The generation of mm10 promoter regions, peak annotation and overlap analyses were performed with R (v4.2.2) packages ChIPseeker (v1.34.1) and TxDb.Mmusculus.UCSC.mm10.knownGene (v3.10.0). Mouse enhancers were downloaded from EnhancerAtlas 2.0 [16] and lifted to mm10 with the UCSC Genome Browser [17] command line tool liftOver. Heatmaps and profile graphs were generated using the plotHeatmap and plotProfile tools of deepTools (v3.5.1). Motif analysis was performed using the MEME-ChIP software.

*Cut&Tag-seq data analysis*

Raw fastq pairs were trimmed using the cutadapt program (v2.7) with the settings "-q 20 -m 25" using the same adapter sequences as described above. Preprocessed reads were aligned to GRCm38 genome using Bowtie2 (v2.3.5) with the settings reported for CUT&Tag [18]. Final reads were retained after removing non-uniquely mapped reads using samtools (v1.9) with "-q 20" [6] and duplicated reads using picard MarkDuplicates (v2.21.4) with "REMOVE_DUPLICATES=true". Peaks were called and browser track files were generated as described above. Differential peaks between two conditions of the same antibody with 2 biological replicates were identified using R package "DiffBind" (v3.8.0). Normalized bigwig signals, heatmaps and profiles were also generated with the deepTools (v3.5.1).

*Gene set enrichment analysis*

GSEA was performed using the 4.1.0 Java desktop application downloaded from GSEA (https://www.gsea-msigdb.org/gsea/index.jsp). For this study, the hallmark gene sets (MSigDB version 7.2) and Kyoto Encyclopedia of Genes and Genomes gene set collections (MSigDB version 7.2) were used in the analysis (http://www.gsea-msigdb.org/ gsea/downloads_archive.jsp). GSEA was performed using the gene set permutation type with the number of permutations set at 1,000. We used a cutoff for the false discovery rate of $P < 0.05$ to identify significantly enriched gene sets.

*Data Availability*

We downloaded ChIP-seq data from GEO website with the following accession numbers: GSE44286 (OCT4, SOX2 and Nanog ChIP-Seq) (Whyte et al. 2013), GSM288354 (KLF4 ChIP-Seq) (Whyte et al. 2013), and GSE113339 (H3K27ac Hi-ChIP) (Di Giammartino et al. 2019).

All sequence data acquired in the work has been submitted to GEO under the following accession numbers:

GSE282372 (RNA-seq, reviewer token, uparkowgpdgjvgx)

GSE282373 (ChIPmentation, reviewer token, wdgzwaoqpbeztyj)

GSE282374 (Cut-&-Tag, reviewer token, qjevgoucbjevpsv)

GSE282375 (ATAC-seq, reviewer token, ajslqkgizlgdvwj)

Figure S1

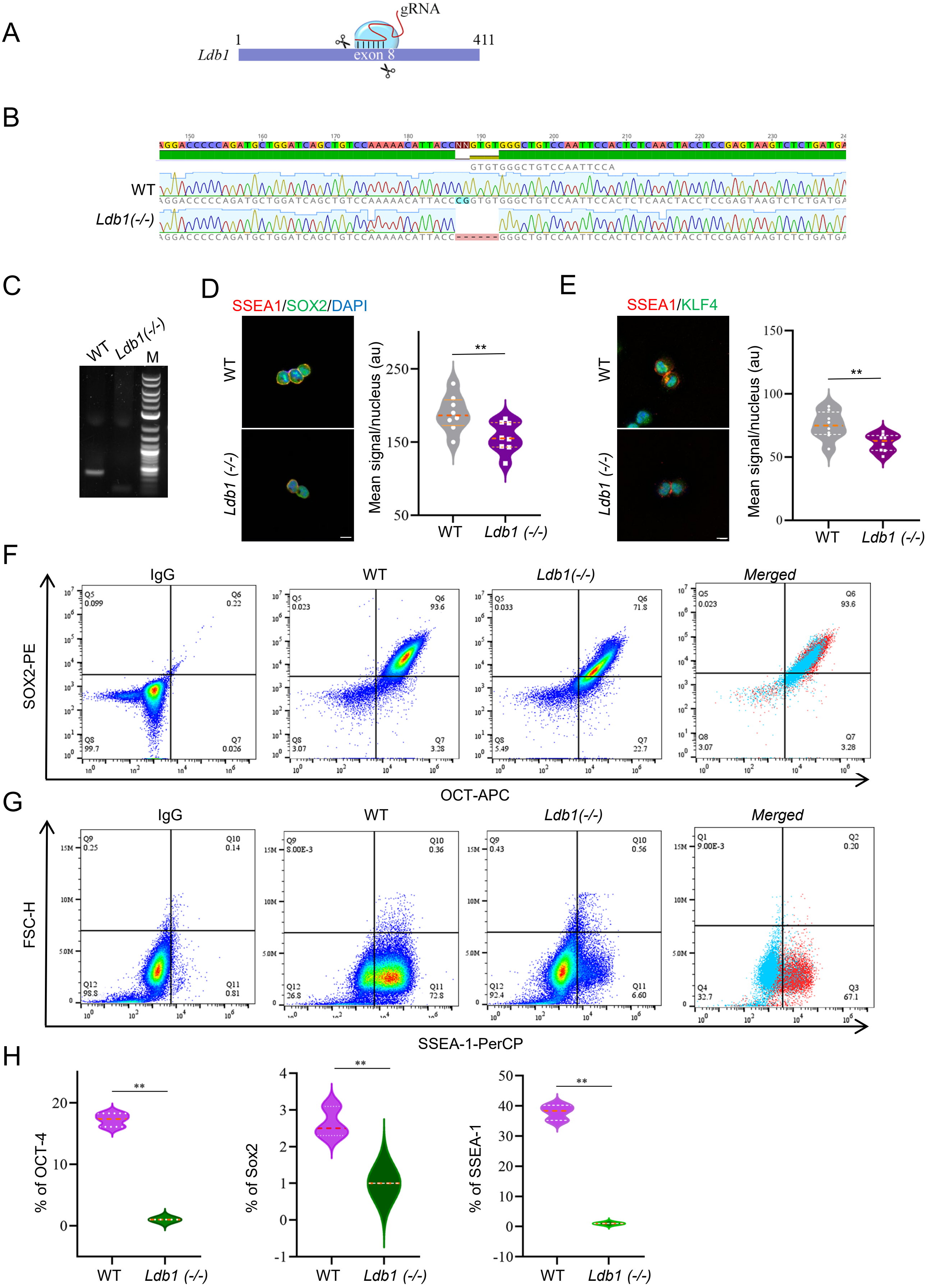
A
gRNA
1
411
Ldb1
exon 8
B
WT
Ldb1(-/-)
C
WT
Ldb1(-/-)
M
D
SSEA1/SOX2/DAPI
WT
Ldb1 (-/-)
Mean signal/nucleus (au)
WT
Ldb1 (-/-)
E
SSEA1/KLF4
WT
Ldb1 (-/-)
Mean signal/nucleus (au)
WT
Ldb1 (-/-)
F
IgG
WT
Ldb1(-/-)
Merged
SOX2-PE
OCT-APC
G
IgG
WT
Ldb1(-/-)
Merged
FSC-H
SSEA-1-PerCP
H
% of OCT-4
% of Sox2
% of SSEA-1
WT
Ldb1 (-/-)

**Figure S1. Pluripotency marker expression is reduced in *Ldb1(-/-)* ESC (Related to Figure 1)**

(A) Schematic representation of the deletion of *Ldb1* exon 8 using CRISPR/cas9 in embryonic stem cells (ESC). (B) Diagram of targeting *Ldb1* using a single guide RNA (sgRNA) at exon 8 generating *Ldb1(-/-)* ESC. (C) PCR analysis demonstrates loss of the targeted locus from genomic DNA of *Ldb1(-/-)* ESC. M, molecular weight markers. (D) Representative immunofluorescence images show the staining of SSEA1 and SOX2 markers with DAPI staining for nuclei in WT and *Ldb1(-/-)* ESC. Right, quantitation of SSEA1 and SOX2 expression levels based on immunofluorescent intensity using ImageJ. The data are mean ± SEM, n=60 cells. (E) Immunofluorescence of KLF4 and DAPI staining for nuclei in WT and *Ldb1(-/-)* ESC. Right, quantitation of KLF4 expression levels based on immunofluorescence intensity using ImageJ. The data are mean ± SEM, n=60 cells. (F) FACS analysis of the expression of SOX2, OCT4, and SSEA1 in WT and *Ldb1(-/-)* ESC. (G) Quantitation of % of cells expressing SOX2, OCT4, and SSEA1 among WT and *Ldb1(-/-)* ESC based on flow cytometry data in panels E and F. The data are mean ± SEM, n=3. **p<0.01. Scale bar, 5 μm.

### Figure S2

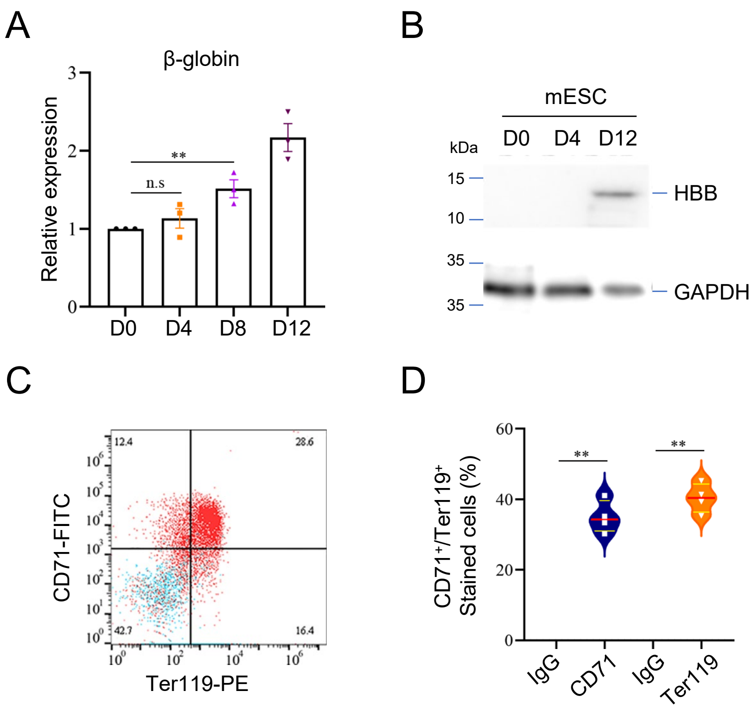


**Figure S2. Appearance of erythroid markers during differentiation of WT EB along the erythroid differentiation pathway (Related to Figure 2)**

(A) Expression of β-globin at different time points (D0, D4, D8, and D12) during differentiation of WT EB. (B) The Western blot shows the protein expression levels of β-globin at different time points during differentiation. (C) Expression of CD71 and Ter119 in the D8 differentiated population of EB. (D) Percentage of cells expressing CD71 and Ter119 in the differentiated population derived from EB at D8 in panel C. The data mean ± SEM, n=3. Statistical significance was determined using a two-tailed Student's t-test, with **, $p<0.01$; "n.s." denotes not significant.

Figure S3

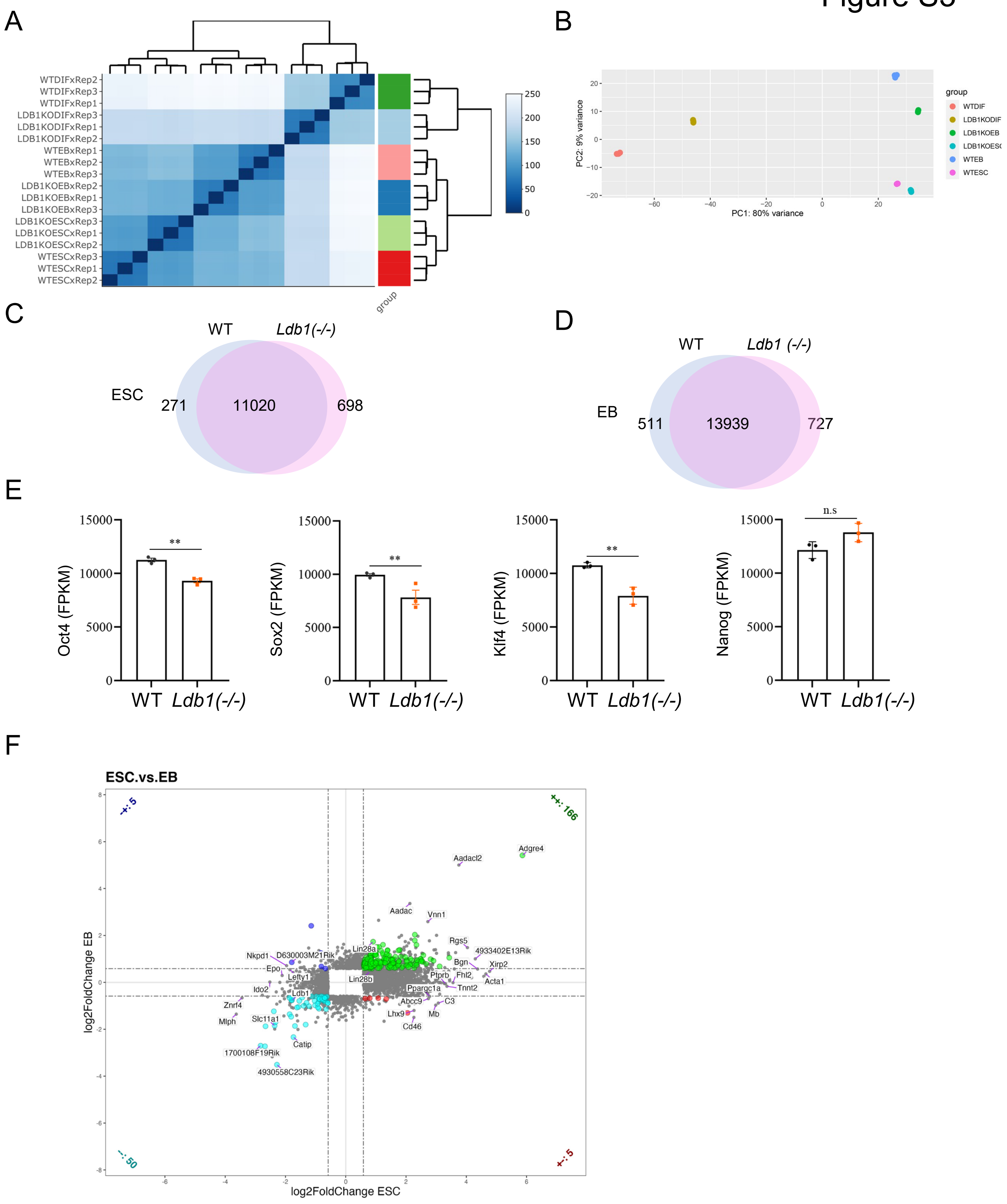
A
WTDIFxRep2
WTDIFxRep3
WTDIFxRep1
LDB1KODIFxRep3
LDB1KODIFxRep1
LDB1KODIFxRep2
WTEBxRep1
WTEBxRep2
WTEBxRep3
LDB1KOEBxRep2
LDB1KOEBxRep1
LDB1KOEBxRep3
LDB1KOESCxRep3
LDB1KOESCxRep1
LDB1KOESCxRep2
WTESCxRep3
WTESCxRep1
WTESCxRep2
group
250
200
150
100
50
0
B
PC1: 80% variance
PC2: 9% variance
group
WTDIF
LDB1KODIF
LDB1KOEB
LDB1KOESC
WTEB
WTESC
C
WT
Ldb1(-/-)
ESC
271
11020
698
D
WT
Ldb1 (-/-)
EB
511
13939
727
E
Oct4 (FPKM)
Sox2 (FPKM)
Klf4 (FPKM)
Nanog (FPKM)
15000
10000
5000
0
**
n.s
WT
Ldb1(-/-)
F
ESC.vs.EB
log2FoldChange ESC
log2FoldChange EB
-+: 5
++: 166
--: 50
+-: 5
Adgre4
Aadacl2
Aadac
Vnn1
Rgs5
4933402E13Rik
Bgn
Xirp2
Ptprb
Fhl2
Acta1
Tnnt2
Ppargc1a
Abcc9
C3
Lhx9
Mb
Cd46
Lin28a
Lin28b
Nkpd1
D630003M21Rik
Epo
Lefty1
Ido2
Ldb1
Znrf4
Mlph
Slc11a1
Catip
1700108F19Rik
4930558C23Rik

**Figure S3. Transcriptomic analysis of WT and *Ldb1*$^{-/-}$ ESC and EB (Related to Figure 3)**

(A) Hierarchical clustering of differential gene expression of WT and *Ldb1(-/-)* ESC, EB, and D8 differentiated samples. (B) Principal components analysis (PCA) plot of WT and *Ldb1(-/-)* ESC, EB, and D8 differentiated cell populations. (C, D) Venn diagrams illustrate the overlap of expressed genes between WT and *Ldb1(-/-)* ESC and EB. (E) FPKM of Oct4, Sox2, Klf4 and Nanog for WT and *Ldb1*$^{-/-}$ ESC from RNA-seq analysis. N=3 biological replicates. (F) Log2fc X log2fc plot of DEGs obtained after LDB1 loss in ESC and EB. Statistical significance determined using a two-tailed Student's t-test, with ** indicating $p<0.01$.

### Figure S4

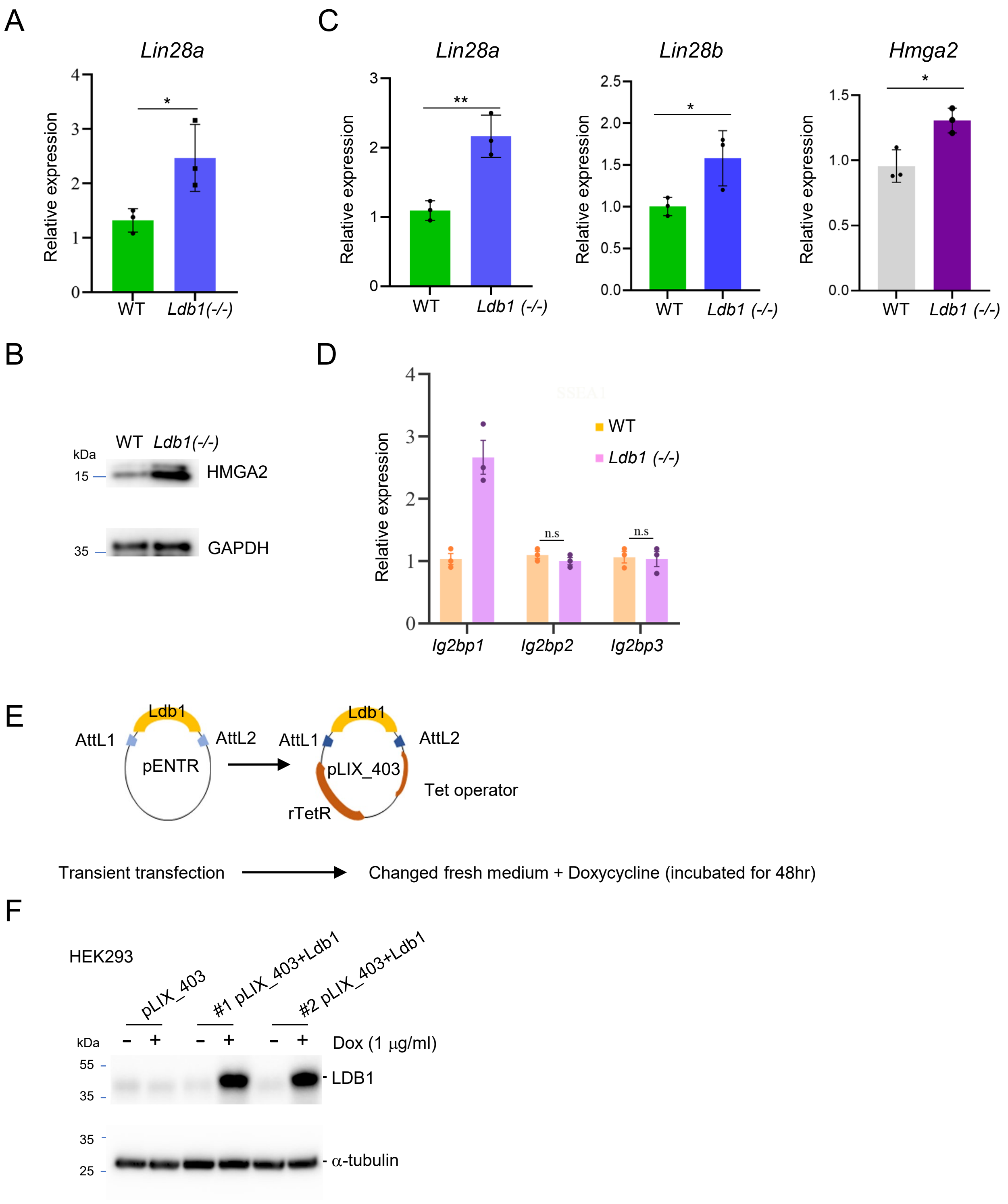
A
Lin28a
Relative expression
WT
Ldb1(-/-)
B
kDa
WT Ldb1(-/-)
HMGA2
GAPDH
C
Lin28a
Lin28b
Hmga2
Relative expression
WT
Ldb1 (-/-)
D
Relative expression
WT
Ldb1 (-/-)
n.s
Ig2bp1
Ig2bp2
Ig2bp3
E
Ldb1
AttL1
AttL2
pENTR
pLIX_403
rTetR
Tet operator
Transient transfection
Changed fresh medium + Doxycycline (incubated for 48hr)
F
HEK293
pLIX_403
#1 pLIX_403+Ldb1
#2 pLIX_403+Ldb1
kDa
Dox (1 μg/ml)
LDB1
α-tubulin

**Figure S4.** ***Let7*** **pathway downstream target genes of LDB1 (Related to Figure 4)** (A) RT-qPCR showing mRNA levels of *Lin28a* in WT and *Ldb1(-/-)* ESC. The data mean ± SEM. N=3 independent experiments. (B) The western blot shows the protein levels of HMGA2 in WT and *Ldb1(-/-)* ESC. (C) RT-qPCR for nascent RNA using intron spanning primers in WT and *Ldb1(-/-) E*SC. (D) RT-qPCR determination of expression of downstream target of *Let7 Ig2bp1*, *Ig2bp2*, and *Ig2bp3* in WT and *Ldb1(-/-)* ESC. (E) Scheme of Tet-inducible LDB1 expression vector (pLIX_403) and transfection into ESC. The vector design allows for controlled expression of LDB1 in response to doxycycline (Dox, 1 μg/ml) induction. (F) Western blot analysis showing the expression levels of LDB1 in HEK293 cells upon doxycycline induction. TUB served as internal control. Statistical significance determined using a two-tailed Student's t-test; **, $p<0.01$.

Figure S5

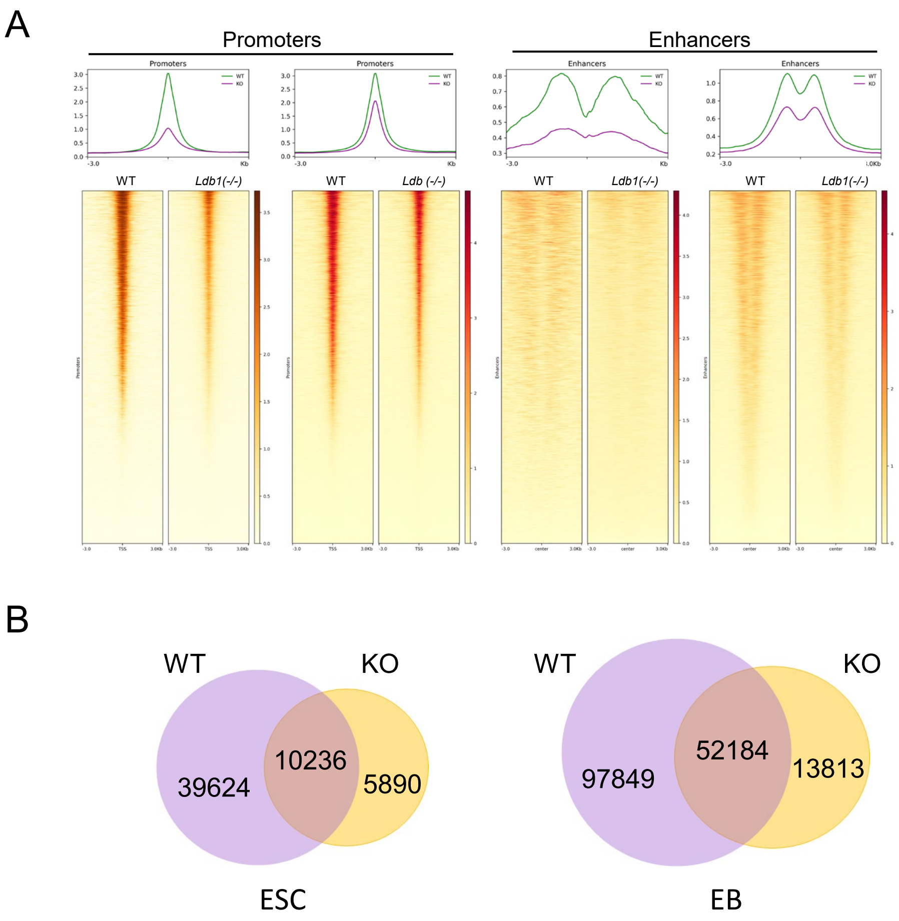


**Figure S5. Chromatin accessibility and histone modification changes is ESC and EB after LDB1 loss (Related to Figure 5)**

(A) Heatmaps show signal intensity for chromatin accessibility in promoters or enhancers in WT and *Ldb1(-/-)* ESC and EB using ATAC-seq. Enhancer Atlas2.0 mm10 (lifted over) was used to determine ATAC-seq peaks at enhancers (ESC_D0). N=2 biological replicates. (B) Venn diagrams showing lost, gained or shared ATAC-seq peaks between WT or *Ldb1(-/-)* ESC or EB.

### Figure S6

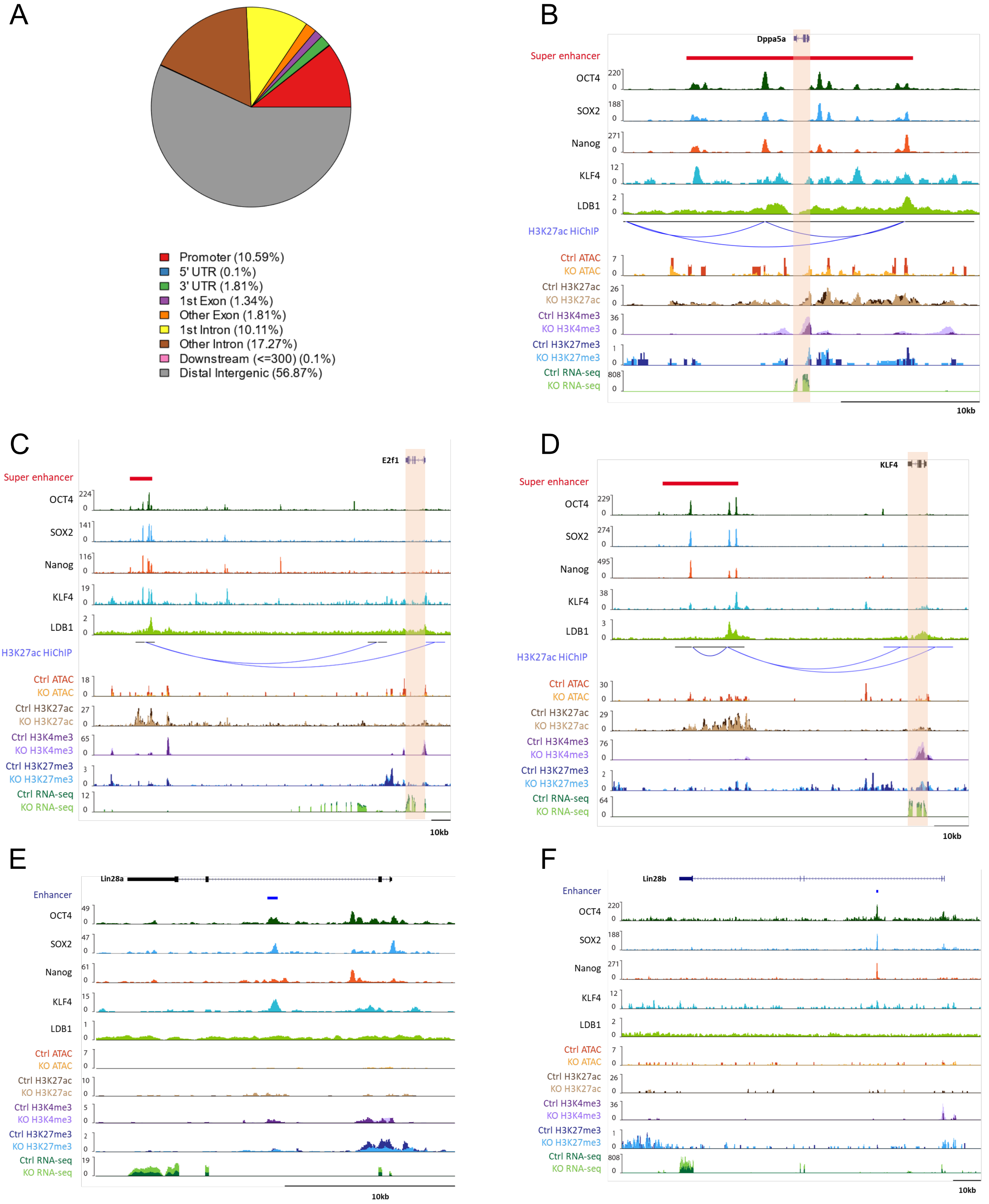

A
Promoter (10.59%)
5' UTR (0.1%)
3' UTR (1.81%)
1st Exon (1.34%)
Other Exon (1.81%)
1st Intron (10.11%)
Other Intron (17.27%)
Downstream (<=300) (0.1%)
Distal Intergenic (56.87%)
B
Dppa5a
Super enhancer
OCT4
SOX2
Nanog
KLF4
LDB1
H3K27ac HiChIP
Ctrl ATAC
KO ATAC
Ctrl H3K27ac
KO H3K27ac
Ctrl H3K4me3
KO H3K4me3
Ctrl H3K27me3
KO H3K27me3
Ctrl RNA-seq
KO RNA-seq
10kb
C
E2f1
Super enhancer
OCT4
SOX2
Nanog
KLF4
LDB1
H3K27ac HiChIP
Ctrl ATAC
KO ATAC
Ctrl H3K27ac
KO H3K27ac
Ctrl H3K4me3
KO H3K4me3
Ctrl H3K27me3
KO H3K27me3
Ctrl RNA-seq
KO RNA-seq
10kb
D
KLF4
Super enhancer
OCT4
SOX2
Nanog
KLF4
LDB1
H3K27ac HiChIP
Ctrl ATAC
KO ATAC
Ctrl H3K27ac
KO H3K27ac
Ctrl H3K4me3
KO H3K4me3
Ctrl H3K27me3
KO H3K27me3
Ctrl RNA-seq
KO RNA-seq
10kb
E
Lin28a
Enhancer
OCT4
SOX2
Nanog
KLF4
LDB1
Ctrl ATAC
KO ATAC
Ctrl H3K27ac
KO H3K27ac
Ctrl H3K4me3
KO H3K4me3
Ctrl H3K27me3
KO H3K27me3
Ctrl RNA-seq
KO RNA-seq
10kb
F
Lin28b
Enhancer
OCT4
SOX2
Nanog
KLF4
LDB1
Ctrl ATAC
KO ATAC
Ctrl H3K27ac
KO H3K27ac
Ctrl H3K4me3
KO H3K4me3
Ctrl H3K27me3
KO H3K27me3
Ctrl RNA-seq
KO RNA-seq
10kb

**Figure S6. Effect of LDB1 loss on transcription and chromatin organization of LDB1 targets (Related to Figure 6)**

(A) Genomic localization of LDB1 peaks in ESC. (B-F) Browser views of transcription factor occupancy and chromatin status at select LDB1 regulated genes (B) *Dppa5a*, (C) *E2f1* and (D) *Klf4,* (E) LIN28a, (F) LIN28b. Data for OCT4, SOX2, Nanog and KLF4 are from [19] and H3K27ac HiChIP is from [20].

Figure S7

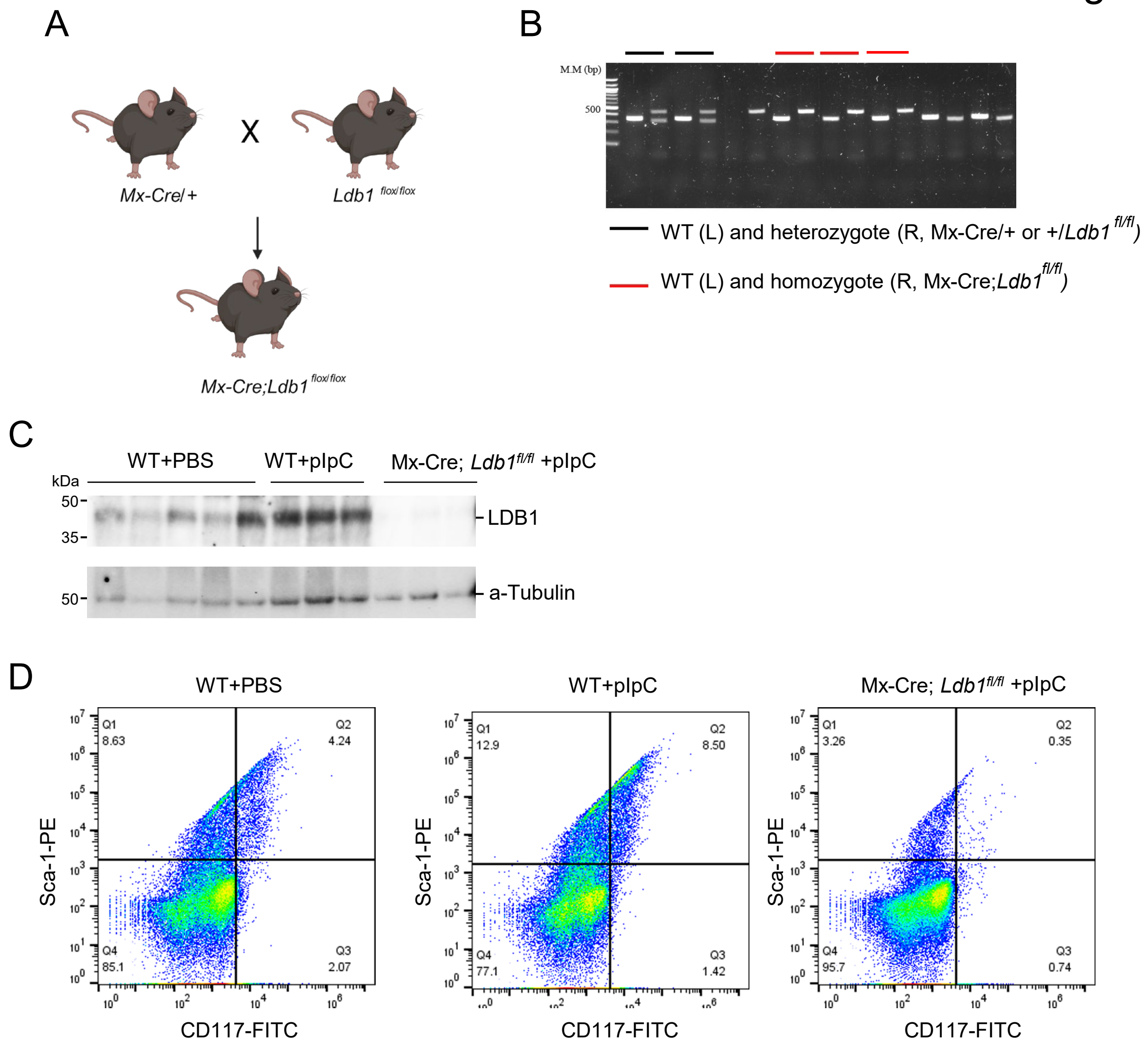


**Figure S7. *Ldb1* conditional knock mice (Related to Figure 7)**

(A) The strategy for generating *Mx-Cre; Ldb1*$^{flox/flox}$ mice is illustrated. (B) Genomic PCR was conducted for genotyping of conditional knockout mice. The red bar indicates homozygous *Mx-Cre; Ldb1*$^{flox/flox}$ mice, while the black and gray bars represent wild-type (WT) and heterozygous mice, respectively. (C) Western blot analysis of LDB1 in isolated liver samples from *Mx-Cre; Ldb1*$^{flox/flox}$ mice. (D) Flow cytometric analysis was performed on isolated bone marrow samples from WT+PBS, WT+pIpC, and *Mx-Cre; Ldb1*$^{flox/flox}$ +pIpC mice using Sca1 and CD117 markers. N=3.

**Supplemental Table S1. Primer used in this study.**

| Name | Forward | Reverse |
|---|---|---|
| **qPCR** | | |
| ***Gapdh*** | GTTGTCTCCTGCGACTTCA | GGTGGTCCAGGGTTTCTTA |
| ***Sox2*** | GAGCTTTGCAGGAAGTTTGC | GCAAGAAGCCTCTCCTTGAA |
| ***Nanog*** | ACCTTGGCTGCCGTCTCTGG | AGCAAAGCCTCCCAATCCCAAACA |
| ***Oct4*** | TTTTGGTACCCCAGGCTATG | GCAGGCACCTCAGTTTGAAT |
| ***Lin28a*** | GGCATCTGTAAGTGGTTCAACG | TGATGATCTAGACCTCCACAGTTGT |
| ***Lin28b*** | CAGACAGGTCACCCCAAGAAG | TTTTGCTCTCCTATTGCTGCA A |
| ***b-globin*** | ATGGTGCACCTGACTGATGCTG | GGTTTAGTGGTACTTGTGAGCC |
| ***Gata-1*** | ATGCCTGTAATCCCAGCACT | TCATGGTGGTAGCTGGTAGC |
| ***Brachury*** | AGGAACCACCGGTCATCG | CGTGTGCGTCAGTGGTGTGTAATG |
| ***Gata-4*** | CTCCTACTCCAGCCCCTACC | GTGGCATTGCTGGAGTTACC |
| ***β-actin*** | ATGAAGATCCTGACCGAGCG | TACTTGCGCTCAGGAGGAG |
| ***Igfbp1*** | GCCCAACAGAAAGCAGGAGATG | GTAGACACACCAGCAGAGTCCA |
| ***Igfbp2*** | CCTCAAGTCAGGCATGAAGGAG | TGGTCCAACTCCTGCTGGCAAG |
| ***Igfbp3*** | CCTCAATGTGCTGAGTCCCAGA | CTTGTCCACACACCAGCAGAAG |
| ***Ldb1*** | AGGAAGCTTTTCCTCTGCAT | CCTGAAGAAACCCAGAGATGA |
| **sgRNA** | | |
| ***Ldb1-1*** | TGGAATTGGACAGCCCACAC | ACGGCTACAGAACTGGACAG |
| ***Lin28a*** | CACCGCACCTTTAAGAAGTCTGCCA | AAACTGGCAGACTTCTTAAAGGTGC |
| ***Ln28b*** | CACCGGAAGTGAAAGAAGACCTAAA | AAACTTTAGGTCTTCTTTCACTTCC |
| ***Ldb1-2*** | CCAAAAACATTACCCGGTGT | GGAGTGTGACAATCTCTGGT |
| **Genotyping PCR** | | |
| ***Cre*** | CGATGCAACGAGTGATGAGG | GACTTGCTGTCACTTGGTCGT |
| ***Ldb1*** | AGGAAGCTTTTCCTCTGCAT | CCTGAAGAAACCCAGAGATGA |

Table S2. Antibodies used in this study

| Reagent or Resource | Source | Identifier | Dilution |
|---|---|---|---|
| **Western blot sections** | | | |
| Rabbit polyclonal anti-mouse-$\alpha$-tubulin | Cell signaling | Cat#77763 | 1:1000 |
| Rabbit polyclonal anti-mouse-$\beta$-actin | Cell signaling | Cat#19069 | 1:1000 |
| Rabbit polyclonal anti-mouse-LDB1 | Cell signaling | Cat#55476 | 1:1000 |
| Rabbi polyclonal anti-mouse-SOX2 | Cell signaling | Cat#3579 | 1:1000 |
| Rabbit polyclonal anti-mouse-NANOG | Abcam | Cat#ab19069 | 1:1000 |
| Rabbit polyclonal anti-mouse-OCT-4 | Abcam | Cat#ab19069 | 1:1000 |
| Rabbit polyclonal anti-mouse-KLF4 | Cell signaling | Cat#12173 | 1:1000 |
| Rabbit polyclonal anti-mouse-HMGA2 | Cell signaling | Cat#8179 | 1:1000 |
| Rabbit polyclonal anti-mouse-LIN28a | Cell signaling | Cat#3695 | 1:1000 |
| Rabbit polyclonal anti-mouse-LIN28b | Cell signaling | Cat#11965 | 1:1000 |
| Rabbit polyclonal anti-mouse-GAPDH | Cell signaling | Cat#5174 | 1:1000 |
| **Immunofluorescence (IF) sections** | | | |
| Rabbit polyclonal Anti-Nanog | Abcam | Cat#ab19069 | 1:200 |
| Mouse monoclonal anti-mouse-SSEA1 | Abcam | Cat#ab19069 | 1:200 |
| Rabbit polyclonal anti-mouse-OCT-4 | Abcam | Cat#ab19069 | 1:200 |
| Rabbi polyclonal anti-mouse-SOX2 | Abcam | Cat#ab3579 | 1:200 |
| **ChIP mentation sections** | | | |
| Rabbit Anti-LDB1 | Abcam | Cat#ab96799 | 1:200 |
| **Cut &Tag** | | | |
| SOX2 antibody (pAb) | Active motif | Cat#39843 | 10 ul |
| OCT-4 antibosy (pAb) | Active motif | Cat#39811 | 10 ul |
| CTCF antibody (pAb) | Active motif | Cat#61311 | 1 ul/50 ul |
| Histone H3K27ac (pAb) | Active motif | Cat#39133 | 1 ul/50 ul |
| Histone H3K4me3 (pAb) | Active motif | Cat#39915 | 1 ul/50 ul |
| Histone H3K27me3 (pAb) | Active motif | Cat#39155 | 1 ug/50 ul |
